# Microtubule-associated IQD9 guides cellulose synthase velocity to shape seed mucilage

**DOI:** 10.1101/2021.12.11.472226

**Authors:** Bo Yang, Gina Stamm, Katharina Bürstenbinder, Cătălin Voiniciuc

**Affiliations:** Independent Junior Research Group–Designer Glycans, Leibniz Institute of Plant Biochemistry, 06120 Halle (Saale), Germany; Department of Molecular Signal Processing, Leibniz Institute of Plant Biochemistry, 06120 Halle (Saale), Germany

**Keywords:** Arabidopsis thaliana, cellulose synthesis, cortical microtubules, matrix polysaccharides, scaffold proteins, seed mucilage, plant cell wall

## Abstract

- Arabidopsis seeds release large capsules of mucilaginous polysaccharides, which are shaped by an intricate network of cellulosic microfibrils. Cellulose synthase complexes is guided by the microtubule cytoskeleton, but it is unclear which proteins mediate this process in the seed coat epidermis (SCE).
- Using reverse genetics, we identified *IQ67 DOMAIN 9* (*IQD9*) and *KINESIN LIGHT CHAIN-RELATED 1* (*KLCR1*) as two highly expressed genes during seed development and comprehensively characterized their roles for cell wall polysaccharide biosynthesis and cortical microtubule (MT) organization.
- Mutations in *IQD9* as well as in *KLCR1* lead to compact mucilage capsules with aberrant cellulose distribution, which can be rescued by transgene complementation. Double mutant analyses revealed that their closest paralogs (*IQD10* and *KLCR2*, respectively) are not required for mucilage biosynthesis. IQD9 physically interacts with KLCR1 and localizes to cortical MTs to maintain their organization in SCE cells. Similar to the previously identified TONNEAU1 (TON1) RECRUITING MOTIF 4 (TRM4) protein, IQD9 is required to maintain the velocity of cellulose synthases.
- Our results demonstrate that IQD9, KLCR1 and TRM4 are MT-associated proteins that are required for seed mucilage architecture. This study provides the first direct evidence that members of the IQD, KLCR and TRM families have overlapping roles in guiding the distribution of cell wall polysaccharides. Therefore, SCE cells provide an attractive system to further decipher the complex genetic regulation of polarized cellulose deposition.

## Introduction

The seed coat epidermal (SCE) cells of some Angiosperms, including the model plant *Arabidopsis thaliana*, synthesize large amounts of hydrophilic polysaccharides (North *et al*., 2014; Voiniciuc *et al*., 2015c; Šola *et al*., 2019). Although the mucilage capsules that rapidly encapsule Arabidopsis seeds upon hydration are pectin-rich, they can be regarded as specialized secondary cell walls because they also contain hemicelluloses that are typical of woody tissues (Voiniciuc *et al*., 2015c). Substituted xylans and heteromannans maintain the attachment of mucilaginous pectin to the seed surface and the organization of ray-like cellulose microfibrils (Yu *et al*., 2014; Voiniciuc *et al*., 2015b,a; Hu *et al*., 2016; Ralet *et al*., 2016). Upon imbibition of a dry seed, expanding mucilage ruptures the outer primary cell wall to release a two-layered gelatinous capsule that can be visualized by ruthenium red (RR), a pectin-binding dye. Cellulosic rays extend from the top of each SCE cell to intertwine and anchor the inner, adherent mucilage layer to the seed surface (Sullivan *et al*., 2011; Mendu *et al*., 2011; Harpaz-Saad *et al*., 2011). However, the genetic factors that modulate the deposition of highly ordered cellulosic structures in seed mucilage remain largely unknown.

The current dogma is that plant crystalline microfibrils are produced by rosette-shaped cellulose synthase (CESA) complexes (CSC) composed of at least three different CESA isoforms and a growing number of interacting proteins (Polko & Kieber, 2019). In Arabidopsis SCE cells, mutations in *CESA3* and *CESA5* have been shown to affect the deposition of cellulose in mucilage pockets. Loss-of-function *cesa5* mutants have a nearly complete loss of adherent mucilage due to reduced cellulose production (Sullivan *et al*., 2011; Mendu *et al*., 2011; Harpaz-Saad *et al*., 2011), while *cesa3* missense mutants lead to milder alterations of mucilage adherence and cellulose organization (Griffiths *et al*., 2015). Several accessory proteins are also known to influence mucilage cellulose synthesis. COBRA-LIKE2 (COBL2) contains a glycosyl-phosphatidylinositol (GPI) anchor and facilitates the assembly of crystalline cellulose by CESA5 (Ben-Tov et al., 2015), while FEI2 (meaning “fat” in Chinese) and SALT-OVERLY SENSITIVE5 (SOS5) mediate pectin adherence to cellulosic rays via an independent mechanism (Griffiths *et al*., 2014, 2016; Ben-Tov *et al*., 2018). CSC assembly and trafficking are maintained by STELLOs (Greek for “to send”; Zhang *et al*., 2016), and negatively regulated by SHOU4 (“thin” in Chinese; Polko *et al*., 2018), identified via a screen for *fei2* suppressors.

Cortical microtubules (MTs) can orient cellulose microfibril deposition by positioning the delivery of CSCs to the plasma membrane (PM) and guiding their subsequent trajectories (Paredez *et al*., 2006; Gutierrez *et al*., 2009; Bringmann *et al*., 2012a). MT-associated proteins, which shape the cytoskeleton in response to environmental or developmental signals (Lloyd & Hussey, 2001; Sedbrook & Kaloriti, 2008), can influence the organization of mucilage polysaccharides. A temperature-sensitive point mutation in *MICROTUBULE ORGANIZATION 1* (*MOR1*) significantly reduced mucilage release at 29°C (McFarlane *et al*., 2008). A second MT-associated protein, TONNEAU1 (TON1) RECRUITING MOTIF 4 (TRM4), was recently found to organize cortical arrays and cellulose distribution (Yang *et al*., 2019). The *trm4* seed mucilage capsules are compact and have shorter cellulosic rays compared to the WT, without altering pectin adherence. Arabidopsis CESAs circle around the cytoplasmic column of SCE cells to polarly deposit cellulose microfibrils in mucilage pockets (Griffiths *et al*., 2015), but the network of proteins that guide CESAs in SCE cells remains unclear.

Plant-specific IQ67 DOMAIN (IQD) proteins associate with MTs and have scaffold-like properties (Bürstenbinder *et al*., 2017). Their eponymous IQ67 domain contains 67 amino acids with calmodulin-recruiting motifs (Abel *et al*., 2005), which could be involved in Ca^2+^ signaling integration (Kölling *et al*., 2019). Since multiple Arabidopsis IQDs control cell shape and size (Bürstenbinder *et al*., 2017; Liang *et al*., 2018; Mitra *et al*., 2019), they are hypothesized to support cell wall deposition. Certain IQDs interact with KINESIN LIGHT CHAIN-RELATEDs (KLCRs; Bürstenbinder et al., 2013; Zang et al., 2021) proteins. KLCRs are also known as CELLULOSE SYNTHASE-MICROTUBULE UNCOUPLING (CMU) and stabilize MTs at the PM during cellulose synthesis (Liu *et al*., 2016). However, direct evidence for how IQDs influence the biosynthesis of cell wall polysaccharides has been lacking.

In this study, we identified IQD9 and KLCR1 as two additional players that maintain MT organization and the deposition of ray-like cellulosic microfibrils in SCE cells. In the absence of *IQD9* or *KLCR1*, mutant seeds released compact mucilage capsules due to aberrant deposition of cellulose microfibrils as previously observed for *trm4* lines. IQD9 physically interacted *in vivo* with KLCR1 and formed filamentous arrays. Live-cell imaging showed that IQD9 and KLCR1 are localized a circular pattern during mucilage biosynthesis, which is reminiscent of CSC trajectories. Both *iqd9* and *trm4* mutants displayed slower CESA3 particles in SCE cells, indicating that these MT-associated proteins guide cellulose deposition. IQD, KLCR and TRM proteins therefore have overlapping roles in cell wall biosynthesis.

## Materials and Methods

### Plant materials and growth conditions

*Arabidopsis thaliana* Col-0 (WT) and T-DNA insertion mutants (Table S1) analyzed in this study were obtained from the Nottingham Arabidopsis Stock Centre, unless otherwise noted. The *proUBQ:RFP-TUB6* in Col-0 (Ambrose *et al*., 2011), *proCESA3:GFP-CESA3* in *je5* mutant (Desprez *et al*., 2007) and *pKLCR1:KLCR1-GFP* in *klcr1-1* (Zang *et al*., 2021) transgenic plants have been described previously. Arabidopsis plants were grown in a phytochamber with constant light (100–120 µmol m^-2^ s^-1^), 22°C and 60% humidity. *Nicotiana benthamiana* plants were grown in a greenhouse (16 h light, 8 h dark) at 22–24°C.

### Arabidopsis transcriptional analyses

Total RNA was isolated from 10-day-old seedlings using TRI Reagent (Sigma Aldrich), according to its manual. The cDNA was prepared from 4 µg of RNA using RevertAid reverse Transcriptase (Thermo Fisher Scientific), and reverse transcription polymerase chain reaction (RT-PCR) was performed using primers listed in Table S2. *ACTIN2* served as a housekeeping gene, and WT genomic DNA was included as a control.

For GUS staining, sample tissues (seedlings, flower buds, siliques) were fixed in 80% (v/v) ice-cold acetone for 30 min and incubated for 4 h to overnight in GUS staining solution (50 mM sodium phosphate, pH 7.2, 0.5 mM K_3_Fe(CN)_6_, 0.5 mM K_4_Fe(CN)_6_, 2 mM 5-bromo-4-chloro-3-indolyl-β-glucuronic acid, 10 mM EDTA) at 37 °C. Images were acquired using a Zeiss Axioplan 2 microscope or a Nikon SMZ 1000. For GUS staining of Arabidopsis seeds, siliques of different ages were opened with forceps at the replum. Seeds were collected with a small spoon and transferred to a 1.5 ml tube with staining solution. Prior to DIC microscopy, staining solution was removed and samples were mounted in chloralhydrate.

### Plasmid construction and plant transformation

Primers used for plasmid construction are listed in Table S2. *Promoter*:*GFP-GUS* constructs were generated by integrating 1.4 kb of the *IQD9* or *IQD10* promoter into pBGWFS7 (Karimi *et al*., 2002). The *pIQD9:IQD9-GFP* transgene was assembled using full-length genomic DNA (from ∼1.5 kb upstream of ATG up to, but excluding, the *IQD9* stop codon) into pB7FWG0 vector. The constructs were stably transformed into Arabidopsis WT plants (for *pIQD:GFP-GUS*) or *iqd9 iqd10* double mutant (for *pIQD9:IQD9-GFP*) respectively via *Agrobacterium*-mediated floral dip transformation. The *p35S:IQD9-GFP, p35S:mCherry-KLCR1*, and *p35S:RFP-KLCR1* transgenes for *N. benthamiana* assays were cloned using Gateway into pB7FWG2 (Karimi *et al*., 2002), pJOG393 (Gantner *et al*., 2018) or pGWB455 (Nakagawa *et al*., 2007) vectors, respectively. The mCherry- and RFP-tagged KLCR1 showed similar results. The *pUBQ10:RFP-TUB6* plasmid for transient expression was previously generated (Yang *et al*., 2019). Transient expression was performed in *N. benthamiana* leaves as previously described (Grefen *et al*., 2010). In short, *Agrobacterium tumefaciens GV3101* cells containing the desired constructs were mixed with the P19 viral suppressor (OD600 = 0.7 for each) and incubated for 4 h (at 18 °C, 200 rpm) before infiltration into the lower side of leaves from 5-week-old plants.

### Staining and quantification of mucilage area

Around 30 seeds were hydrated in water for 30 min and stained with 300 µl of 0.01% (w/v) RR (Sigma-Aldrich; R2751) for 15 min at 125 rpm in 24-well plates. After rinsing with water, the stained seeds were re-suspended in 300 µl of water and imaged with a Leica M165FC stereomicroscope equipped with MC170 HD camera. The mucilage and seed projected areas were quantified using an existing ImageJ pipeline (Voiniciuc *et al*., 2015b).

Cellulose around hydrated seeds was stained with 0.01% (w/v) Pontamine fast scarlet 4B (S4B, also known as Direct Red 23 [Sigma-Aldrich; 212490]) in 50 mM NaCl for 60 min at 125 rpm in 24-well plates (Anderson *et al*., 2010; Mendu *et al*., 2011). The counterstain was performed by mixing with 25 ug mL^-1^ Calcofluor (Megazyme C-CLFR) for 5 min. Seeds were imaged with a Carl Zeiss LSM 780 microscope with 10X/0.45 objective and the following excitation / emission wavelengths (S4B: 561 / 580–650 nm; Calcofluor: 405 / 410– 452 nm). The lengths of cellulosic rays were measured by ImageJ.

To view surface morphology, around 30 seeds were mixed with 500 µL of 0.01% (w/v) propidium iodide for 15 min. Seeds were rinsed twice with water and imaged using a Leica LSM 900 with 10X/0.3 objective (excitation 488 nm, emission 600-650 nm).

### Seed polysaccharide quantification

Non-adherent mucilage was extracted by gently mixing 5 mg seeds in water for 30 min at 125 rpm and subsequently the adherent mucilage was isolated using a ball mill (Retsch; MM400) for 30 min at 30 Hz, as previously described (Voiniciuc, 2016). The two mucilage fractions, spiked with ribose and inositol respectively, were hydrolyzed and quantified via high-performance anion exchange chromatography with pulsed amperometric detection (HPAEC-PAD), as described (Voiniciuc & Günl, 2016) with the following changes. HPAEC-PAD was performed on a Metrohm 940 Professional IC Vario, using Metrosep Carb 2-250/4.0 columns and a published gradient (Mielke *et al*., 2021). Peaks were integrated and calibrated (manually corrected if necessary) in the MagIC Net 3.2 software (Metrohm).

Crystalline cellulose was quantified using the Updegraff reagent (Updegraff, 1969) and the anthrone colorimetric assay (Foster *et al*., 2010), as previously adapted for Arabidopsis whole seeds (Voiniciuc *et al*., 2015b).

### Salt stress treatments

Germination assays were performed in 24-well culture plates as described previously (Yang et al., 2021). Around 35 seeds were hydrated in 500 µl of water or 150 mM CaCl_2_ solution per well. All the seeds were vernalized for 66 h (dark, 4°C), transferred to a chamber with constant light (100– 120 µmol m^-2^ s^-1^), 22°C and 60% humidity. The seeds were imaged every 24 h with a Leica M165FC stereomicroscope and defined as germinated when the radicle length was > 70 µm.

For the seedling salt stress assay, the seeds were placed on ½ Murashige and Skoog agar plates, stratified for 66 h (dark, 4°C) and grown vertically in the climate-controlled chamber described above. Five-day-old seedlings of similar size were transferred to fresh agar plates with or without 100 mM NaCl and growth was imaged using a Nikon D5600 digital camera.

### Stem cell wall analyses

Stem sections were cut by hand from the basal third of stems of 6-week-old plants and were stained in 0.01% toluidine blue O (Sigma-Aldrich; T3260) for 2 min. Stained sections were rinsed twice with water and imaged on a Axioplan 2 with Axiovision software (Zeiss).

The bottom 7 cm of mature stems were harvested and homogenized using a ball mill (Retsch MM400) for 10 min at 30 Hz. The AIR was extracted by sequential washes with 1 mL of 70% (v/v) ethanol, 1 mL of 1:1 (v/v) chloroform:methanol and 1 mL of acetone. Stem AIR was hydrolyzed and quantified via HPAEC-PAD exactly as described for seed mucilage, using ribose as the internal standard.

### Confocal microscopy and image analysis

For fluorescence co-localization assays in *N. benthamiana*, leaf discs at 3 days post-infiltration were imaged with a Zeiss LSM880 inverted confocal microscope using a 40X/1.2 water-immersion objective. For oryzalin treatment, small pieces of transfected leaf were immersed with 1% (v/v) dimethylsulfoxide (DMSO) as mock treatment or 1% (v/v) DMSO containing 0.1 mM oryzalin (Sigma-Aldrich) for 4 h.

For the fluorescence imaging of the Arabidopsis SCE cells, seeds were carefully dissected from siliques at specific stages and were imaged using a Zeiss LSM880 in Airyscan mode with a 40X/1.2 water-immersion objective. For PM, 7 DPA (days post-anthesis) seeds were pre-stained in 50 µM FM4-64 for 30 min before imaging. The subcellular localization of GFP-CESA3 was detected using a 63X/1.4 oil-immersion objective and Airyscan mode, with 330 ms exposure time based on previously described protocols (Vellosillo *et al*., 2015; Duncombe *et al*., 2020). Unless stated otherwise, the time-lapse series were acquired every 5 sec for 5 min. All the samples were mounted on confocal dishes with spacers (VWR International; 734-2905) and were examined with the following excitation/emission settings: GFP (488 / 505–530 nm), RFP/mCherry (514 / 580–635 nm), FM4-64 (514 / 600–700 nm).

For hypocotyl imaging, seeds were sowed on ½ MS agar plates and stratified for 66 h (dark, 4°C). The plates were exposure to light for 6 h at room temperature and then were wrapped with aluminum foil to keep the plates in the dark at room temperature. For the observation of RFP-TUB6 in the hypocotyl epidermal cells, the inner face of epidermal cells in zone 1 of 4-d-old dark-grown seedlings were examined as described previously (Crowell et al., 2011).

All images were processed uniformly using ImageJ. The maximum projection of the Z-stack or time-lapse view was generated by frames using the Z Project tool and average intensity. For the colocalization evaluation, the intensity plot analysis was done by “RGB Profile Plot” plugin. The Pearson correlation coefficient of region of interest (ROI) from single frame was quantified with “Coloc 2”. Kymograph analysis of proteins and velocity quantification of GFP-CESA3 were performed as previously described (Vellosillo *et al*., 2015). Briefly, the time-lapse stack was generated with “Walking Average” plugin. The GFP-CESA3 track was depicted using segmented line on time average image and transferred to time-lapse stack. The kymographs were generated using the “MultipleKymograph” plugin, and the slope of each line was used to calculate the particle velocity.

### Protein-Protein Interactions

Proteins were extracted from transiently transformed *N. benthamiana* leaves using 1 ml lysis buffer (20 mM HEPES, pH 7.5, 40 mM KCl, 1 mM EDTA, 0.1% [v/v] Triton X-100, 1× protease inhibitor cocktail, 1 mM phenylmethylsulfonyl fluoride, and 10% [v/v] glycerol) (Ganguly *et al*., 2020). Homogenate from 500 mg of plant material was centrifuged twice at 4°C at 15000 g for 10 min, and 900 µl of the supernatant was incubated with 20 µl GFP-Traps (Chromotek) overnight at 4°C on a rocking shaker. Beads were equilibrated following manufacturer’s manual using lysis buffer, without Triton-X. The next day, beads were washed four times and were boiled in 80 µl 2x Laemmli Buffer. For each sample, 40 µl were loaded on a SDS gel and blotted afterwards for 1 h. For protein detection, 1:1000 dilution of the GFP antibody 3H9 (Chromotek) or the RFP antibody 6G6 (Chromotek) was used. Anti-mouse (Sigma A9044 1:20000) and anti-rat (Thermo 31470 1:3000) secondary antibodies conjugated to horseradish peroxidase were used to detect RFP and GFP signals, respectively. Western blot images were acquired with a FluorChem system, using 1:1 mixture of Amersham ECL-Prime and ECL-Select as chemiluminescent detection reagents.

## Results

### *IQD9* and *KLCR1* control seed mucilage architecture

We hypothesized that one or more *IQD* genes are involved in Arabidopsis seed mucilage biosynthesis and screened their expression profiles using published transcriptional datasets. *IQD9* (At2G33990) and *IQD10* (At3G15050), its closest paralog, were upregulated in the seed coat at the stages of mucilage biosynthesis (Fig. 1a; Winter *et al*., 2007; Le *et al*., 2010). T-DNA insertions in these genes disrupted their transcription (Fig. 1b) and were screened for seed mucilage defects. While *iqd10* mutant seeds resembled WT, two independent *iqd9* mutants displayed compact RR-stained mucilage capsules (Fig. 1c). Consistent with microarray data and the seed mucilage defects (Fig. 1), *promoter*:GUS reporter assays only showed high seed coat activity for *pIQD9* (Fig. S1). *IQD9* and *IQD10* displayed partially overlapping expression profiles but share only 47% amino acid identity (Fig. S1). Furthermore, *iqd9-1 iqd10-1* (*i9 i10*) double mutant seeds showed compact mucilage capsules resembling the *iqd9* single mutants (Fig. 1c).

**Fig. 1.**
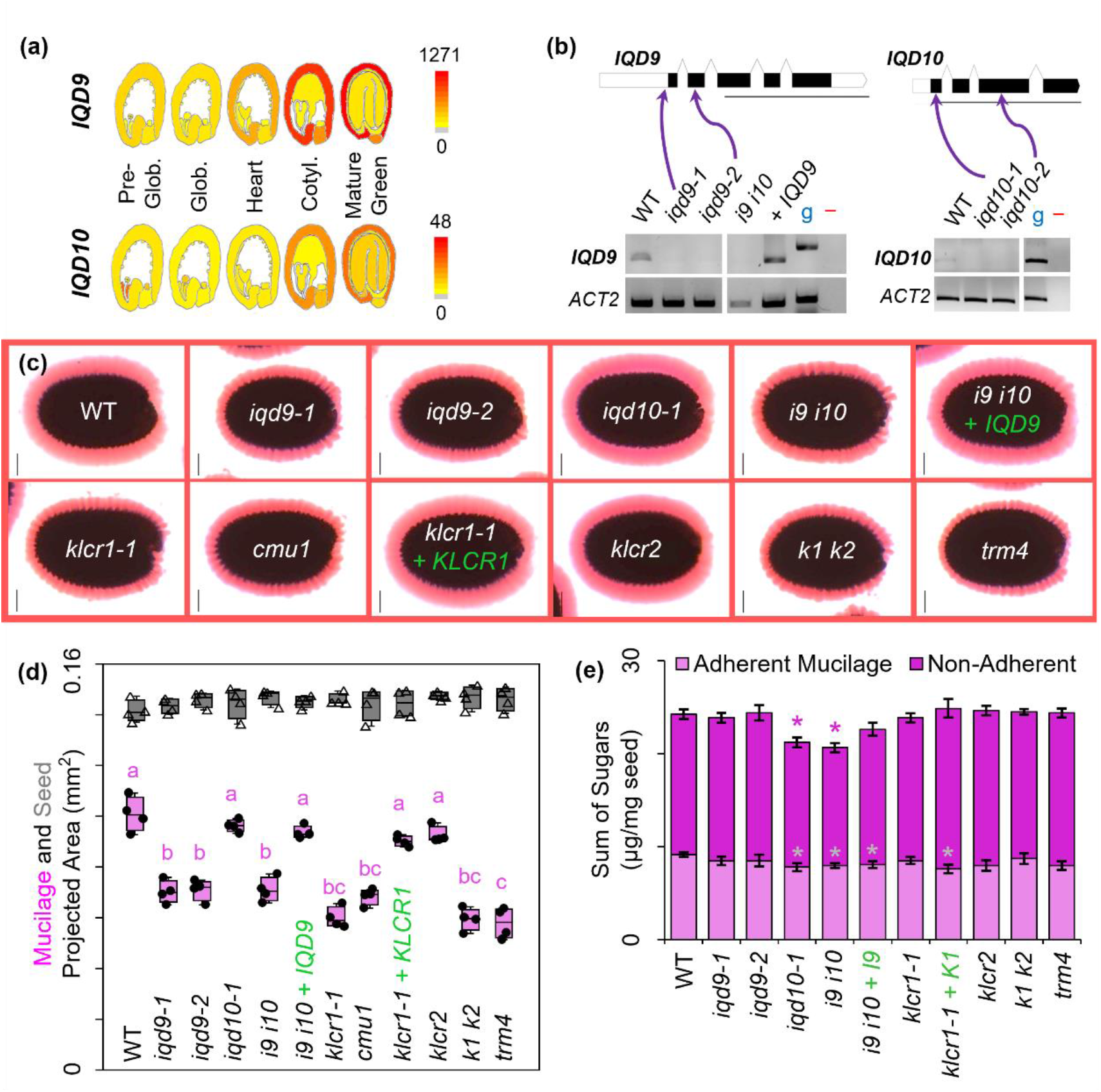
Mutation of *IQD9* and *KLCR1* caused compact mucilage capsule. (a) Expression profiles in the seed eFP browser (Winter *et al*., 2007; Le *et al*., 2010), including absolute expression values. Glob. (globular); Cotyl. (cotyledon). (b) UTR, intron and exon structure of candidate genes. The position and the effects of T-DNA insertions were verified using RT-PCR, *ACTIN2* as a reference gene, g as genomic DNA control, and – as no DNA control. Transgene complemented were marked by + *IQD* or + *KLCR*. Scale bars for gene models = 1000 bp. (c) RR staining of adherent mucilage capsules after gentle shaking in water. Bars = 100 µm. (d) Seed (triangles) and RR-stained mucilage (black dots) area of four biological replicates (>20 seeds each) per genotype. Boxes show the 25–75% quartiles, the median value (inner horizontal line), and whiskers extending to the largest/smallest values. Different letters mark *P* < 0.01 for one-way ANOVA with Tukey test. (e) Absolute amounts of monosaccharides in sequentially-extracted mucilage fractions. Data show mean ±SD of 5 biological replicates, and asterisks mark significant changes compared to WT (Student’s t-test, *P* < 0.001). See Fig. 2 for detailed composition.

Since IQD-KLCR interactions have been reported (Bürstenbinder *et al*., 2013), we also assessed if *KLCR1*/*CMU1* (At4g10840) and *KLCR2*/*CMU2* (At3g27960) are involved in mucilage biosynthesis. *KLCR1* was highly expressed throughout the seed development, while *KLCR2* transcription peaks at the pre-globular and globular stages, before mucilage biosynthesis (Fig. S2a). Two knockout *klcr1* alleles, *klcr1-1* and *cmu1* (Fig. S2b, Table S1), resembled the *iqd9* compact mucilage defect (Fig. 1a), while *klcr2* seeds displayed WT-like mucilage. Both *iqd9* as well as *klcr1* mutants reduced mucilage capsule area by 30–40% compared to WT (Fig. 1d), without altering seed size. The double mutant *i9 i10* and *klcr1-1 klcr2-2* (*k1 k2*) phenocopied the mucilage structure of *iqd9-1* and *klcr1-1* respectively, indicating no functional redundancy between the related genes. Transgene complementation of *i9 i10* with *IQD9* and of *klcr1-1* with *KLCR1* fully rescued the compact mucilage defects (Fig. 1). Both *iqd9* and *klcr1* mutants resembled the mucilage phenotype of *trm4*, which has SCE cells with disorganized MTs (Yang *et al*., 2019). Moreover, *iqd9 klcr1* and *iqd9 trm4* double mutants displayed compact RR-stained mucilage capsules equivalent to the single mutants (Fig. S2c–f).

### *IQD9* and *KLCR1* are specifically required for cellulose distribution in mucilage

Monosaccharide analysis of non-adherent and adherent mucilage fractions revealed that the *iqd9, klcr1* and *trm4* mutants did not alter the content or adherence of matrix polysaccharides to the seed surface (Fig. 1e). Surprisingly, despite no impact on RR staining, the *iqd10-1* mutation correlated with a small but statistically significant reduction in total extractable monosaccharides (Fig. 1c–e). The *iqd10-1* mutation decreased rhamnose (Rha) and galacturonic acid (GalA) content in non-adherent mucilage, along with galactose (Gal) and arabinose (Ara) in adherent mucilage (Fig. 2). Nevertheless, compared to the galactoglucomannan-deficient *muci10* mutant with compact mucilage, all the MT-related mutant seeds released relatively normal levels of mucilage glycans (Fig. 2).

**Fig. 2.**
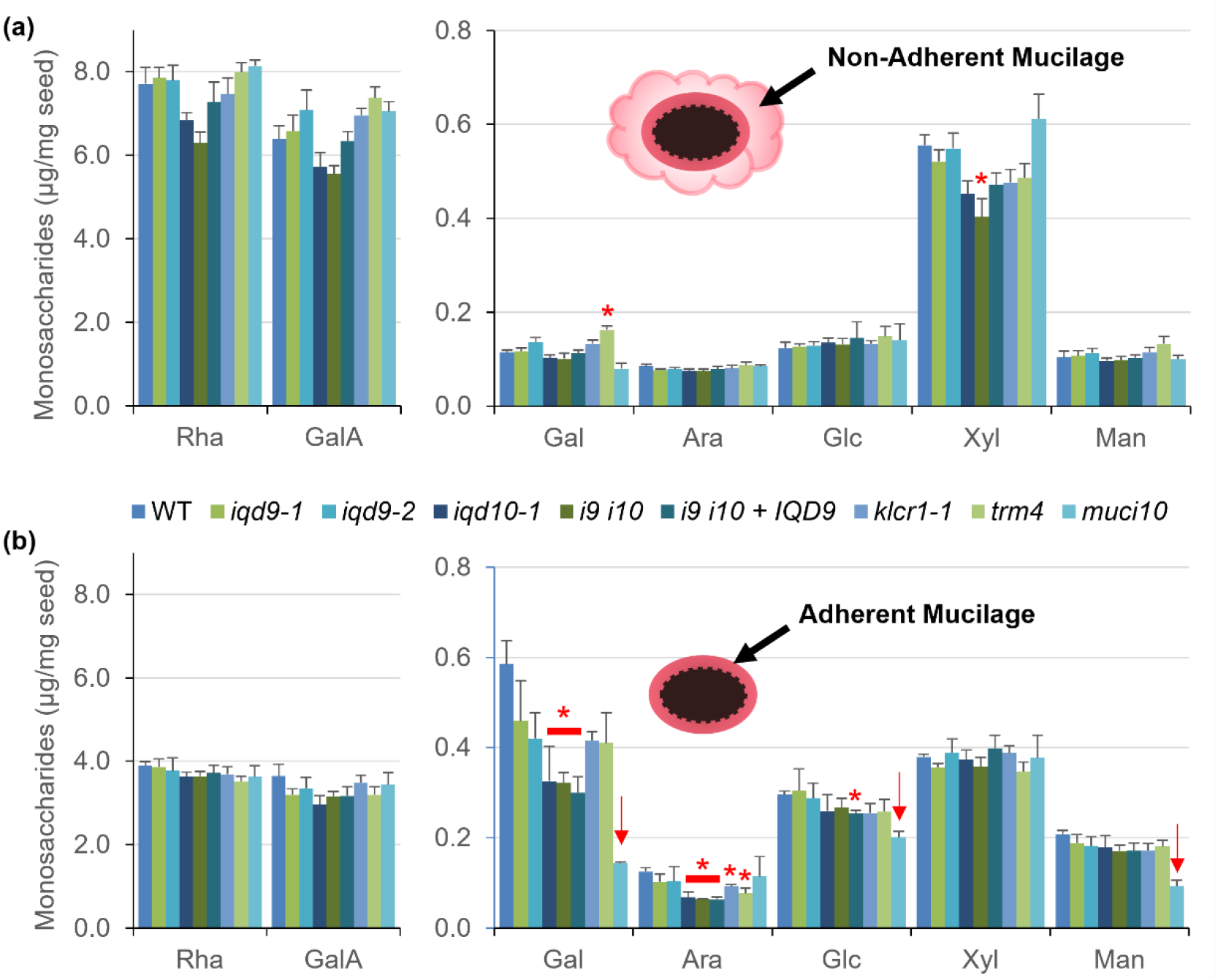
Composition of matrix polysaccharides in seed mucilage extracts. (a) Non-adherent and (b) adherent mucilage polysaccharides were sequentially extracted using water and different mixing intensities. Data show mean + SD of 5 biological replicates (only 3 for *muci10*). Red asterisks and arrows (for galactoglucomannan subunits) mark differences from WT (Student’s t-test, *P* < 0.0001).

Since the matrix polysaccharide composition could not account for the compact mucilage defects of *iqd9* and *klcr1*, we then examined the structure of cellulose using S4B, a specific fluorescent dye (Anderson et al., 2010). The *iqd9* and *klcr1* mutant seeds extruded less cellulose upon hydration (Fig. 3a) and had ∼30% shorter rays atop each columella compared with WT (Fig. 3b). Moreover, the mutant seeds lacked the diffuse cellulose staining that was observed between the WT rays. The S4B-stained seeds of *i9 i10* and *k1 k2* resembled the *iqd9* and *klcr1* single mutants, while *iqd10* and *klcr2* had WT-like seeds. The cellulose defects of *i9 i10* and *klcr1* mutants were rescued by *IQD9* and *KLCR1* transgene complementation, respectively, using their native promoters (Fig. 3a). While all the *iqd9, klcr1* and *trm4* mutant combinations examined showed short S4B-stained cellulosic rays compared to WT (Fig. 3 and Fig. S3), counterstaining of mucilage with calcofluor displayed relatively normal content of other β-glucans (Fig. S3). Despite evident changes in the architecture of cellulose extruded from hydrated seeds, these MT-related mutants did not alter the crystalline cellulose content of whole seeds (Fig. 3c). Mutations in *IQD9* or *KLCR1* did not alter seed shape (Fig. S4), nor the germination rate or sensitivity to salt (Fig. S5).

**Fig. 3.**
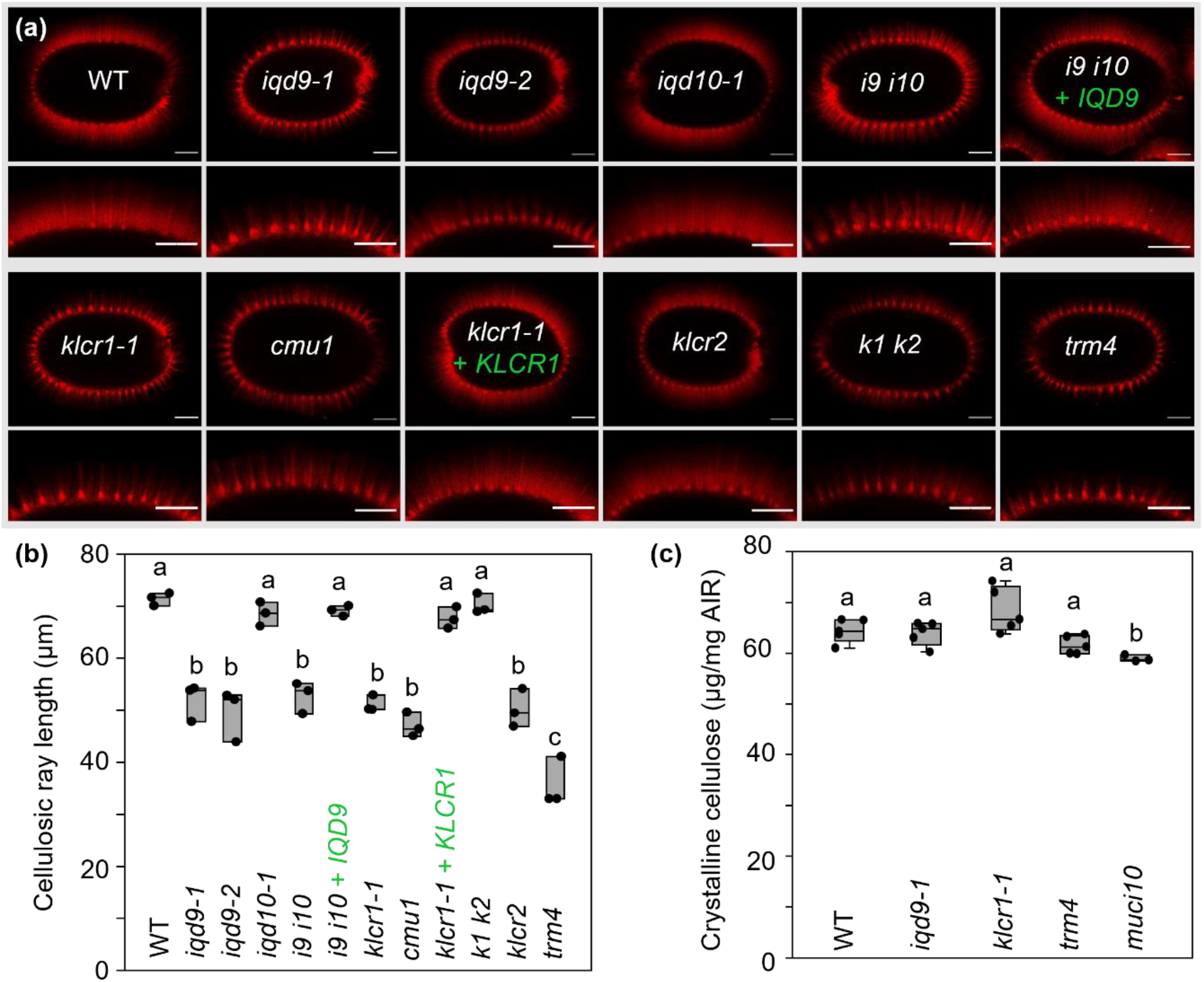
*IQD9* and *KLCR1* are important for cellulose deposition around seed surface. (a) S4B-stained cellulosic rays in mucilage capsules. Bars = 100 µm. (b) The length of cellulosic rays stained with S4B. Boxes show the 25–75% quartiles, the median value (inner horizontal line), and whiskers extending to the largest/smallest values (≥10 measurements per biological replicate). (c) Crystalline cellulose content in whole seeds (5 biological replicates per genotype, except 3 for *muci10*). Different letters in (b) and (c) mark significant changes (one-way ANOVA with Tukey test, P < 0.01).

Therefore, like *TRM4* (Yang *et al*., 2019), the expression of *IQD9* and *KLCR1* in seeds primarily affect mucilage polymer organization.

### *IQD9* and *IQD10* are also expressed beyond the seed coat

Based on *promoter*:GUS reporter fusions, *pIQD9* and *pIQD10* promoters were active in multiple plant tissues, including the vasculature system (shoot and roots), developing flowers and siliques (Fig. S1). Except for anthers, *pIQD9* showed higher activity than *pIQD10* in reproductive organs and seeds (Fig. S2). *IQD10* was expressed highest in Arabidopsis stems and its ortholog in *Populus deltoides* (*PdIQD10*) affects the development of the woody stem (Badmi *et al*., 2018). However, stem cross-sections of the *iqd10* single mutants and the *i9 i10* double mutant did not show the *irregular xylem* (*irx*) phenotype observed in secondary cell wall mutants such as *irx14* (Fig. S6a). Furthermore, *iqd* stems had normal monosaccharide composition (Fig. S6b), while *irx14* stems were xylan-deficient as previously described (Brown *et al*., 2007). *IQD9* and *IQD10* are thus not indispensable for the formation of xylem cells with thick secondary cell walls, or their functions could be masked by other IQDs.

### IQD9 proteins associate with MTs and interact with KLCR1

To determine the subcellular localization of IQD9 proteins, we first co-expressed IQD9-GFP fusion proteins and the MT marker RFP-TUB6 in *N. benthamiana* leaf epidermal cells. IQD9-GFP localized in striated arrays that overlapped with RFP-TUB6 at the cell cortex (Fig. S7), and could be abolished by treating cells with MT-depolymerizing oryzalin. Next, we co-expressed IQD9-GFP with mCherry-tagged KLCR1 (mCherry-KLCR1) in *N. benthamiana* and found that they were co-localized in arrays resembling MTs (Fig. 4a–c). IQD9 and KLCR1 still co-localized when their striated patterns were disassembled by oryzalin treatment (Fig. 4c–e). We validated that these two proteins physically interact using co-immunoprecipitation (co-IP). IQD9-GFP and RFP-tagged KLCR1 (RFP-KLCR1) proteins were expressed in *N. benthamiana* leaves, extracted and purified by GFP-trap beads. Western blotting showed that all recombinant proteins were present in the input fractions (Fig. 4f), before the addition of the GFP-binding beads. RFP-KLCR1 proteins were detected on GFP-Trap beads only when co-expressed with IQD9-GFP, but not with GFP alone. Therefore, IQD9 proteins physically interacted with KLCR1 *in vivo* and were closely associated with cortical MTs.

**Fig. 4.**
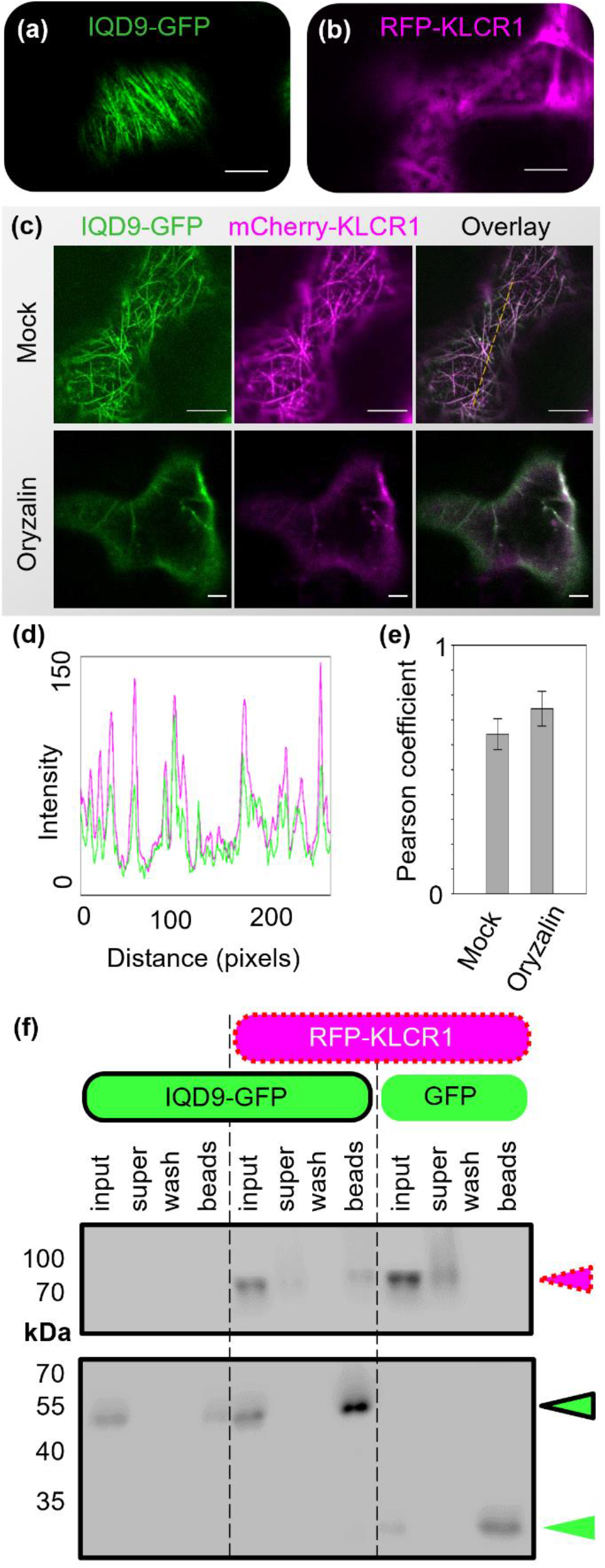
IQD9 co-aligns and interacts with KLCR1. (a) IQD9-GFP localized in cortical arrays in *N. benthamiana* cells. (b) RFP-KLCR1 shows diffuse localization when overexpressed on its own in tobacco. (c) Subcellular co-localization of IQD9-GFP and mCherry-KLCR1 in the mock and oryzalin-treated tobacco epidermal cells. Both transiently expressed proteins were oryzalin-sensitive. (d) Fluorescent intensity plot along the dashed line in (c). Bars = 10 µm. (e) Pearson correlation coefficient between IQD9-GFP and mCherry-KLCR1 in (c), n=5 cells from 5 independent treatments. (f) Co-IP of proteins transiently expressed in tobacco leaves. Colored triangles marked the expected size of each protein. Labels: Input (protein supernatant before adding the GFP-Trap beads); Super (unbound supernatant after bead incubation); Wash (Supernatant from last wash step); Beads (co-IP proteins that tightly bind GFP-Trap).

### Localization of IQD9-GFP during mucilage biosynthesis

To investigate the distribution of IQD9-GFP in Arabidopsis, we examined its subcellular localization under the control of its native promoter in the complemented *iqd9* line, which rescued the mucilage defects (Fig. 1). While undetectable in young seedlings, IQD9-GFP fluorescence was evident during seed coat development (Fig. S8a), particularly at the peak stage of mucilage biosynthesis. Z-stack maximum projections displayed IQD9-GFP proteins in MT arrays, near the PM and inside the nucleus (Fig. 5a). IQD9-GFP displayed circular arrays around the cytoplasmic column, resembling previously described CESA trajectories during mucilage production (Griffiths *et al*., 2015). At SCE cell boundaries, IQD9-GFP proteins co-localized with the PM stained by FM4-64. Time-lapse imaging revealed that IQD9-GFP proteins were static (the vertical lines in kymograph; Fig. 5b), as previously noted for KLCR/CMU proteins (Liu *et al*., 2016). Highly immobile KLCR1-GFP proteins, expressed under its native promoter in the complemented *klcr1* line, were also associated with both MTs and PM throughout SCE development (Fig. S8b), but lacked the nuclear localization observed for IQD9-GFP. In cross-sectional views of live SCE cells, both IQD9-GFP and KLCR1-GFP were localized primarily as striated arrays adjacent to the mucilage pocket (Fig. 5c,d).

**Fig. 5.**
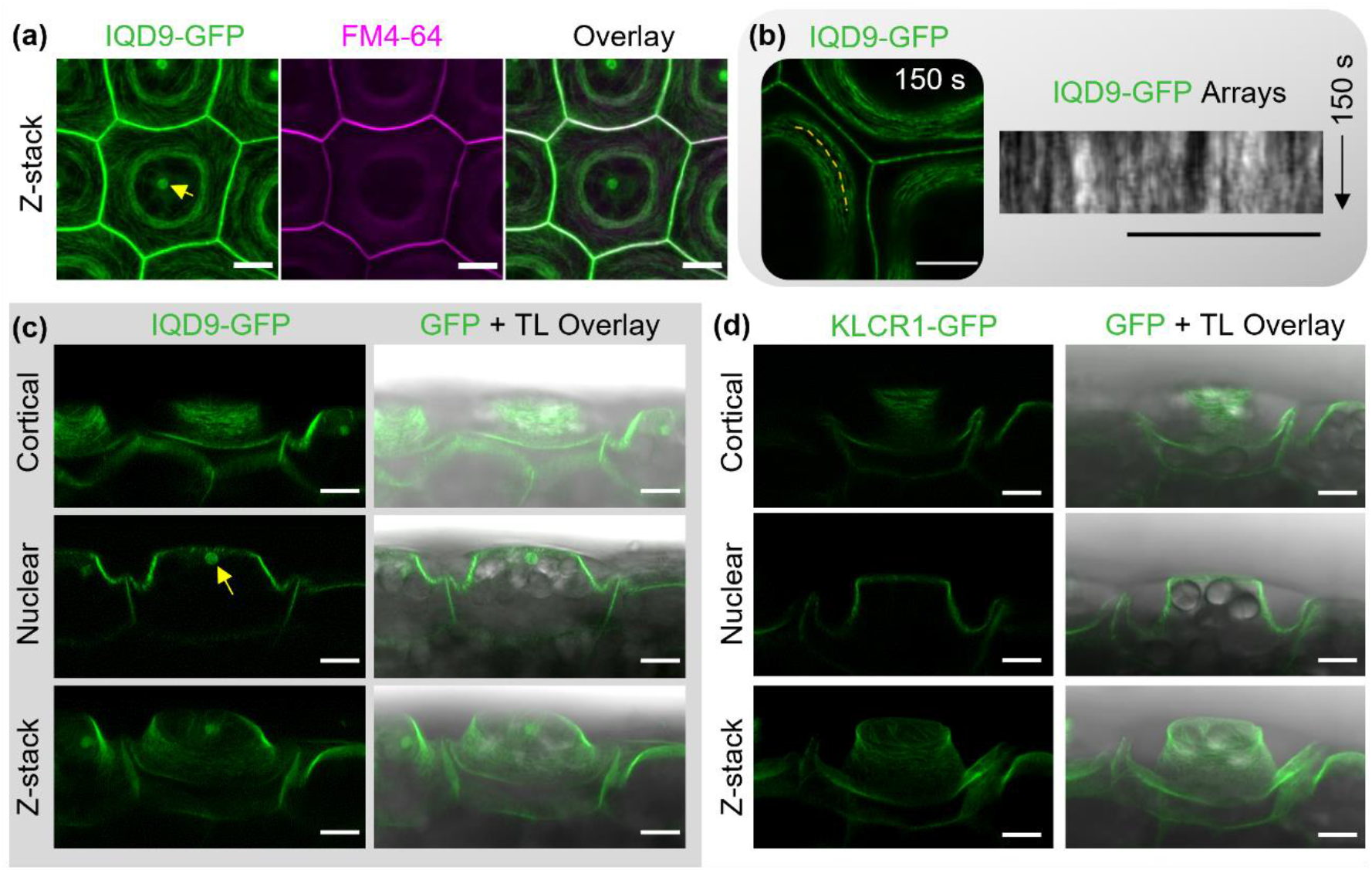
IQD9 and KLCR1 localization during mucilage biosynthesis in complemented lines. (a) Z-stack maximum projection of IQD9-GFP SCE cells stained with FM4-64 at 7 DPA. IQD9 is localized at the PM, MTs and in a nuclear body (arrow). (b) Time-lapse of IQD9-GFP and kymograph along the dashed line. (c) Cross-sectional views of SCE cells expressing IQD9-GFP. The arrow marks a nuclear compartment. (d) Cross-sectional views of SCE cells expressing KLCR1-GFP during mucilage biosynthesis. Bars = 10 µm.

### *IQD9* maintains MT organization in SCE cells

Proper MT organization is essential for the establishment of mucilage architecture. The MT marker RFP-TUB6, which formed circular arrays around the cytoplasmic column of SCE cells at 7 DPA (Yang *et al*., 2019), was introduced into the *iqd9-1* mutant by crossing. In contrast to the WT background, circular MT arrays were undetectable in all *iqd9* SCE cells expressing RFP-TUB6 (Fig. 6). While MT organization was severely disrupted in the seed coat, both WT and *iqd9* displayed transversely oriented RFP-TUB6 arrays in hypocotyl epidermal cells (Fig. S9), despite some variation in fluorescence intensity. Therefore, the distribution of cortical MTs in the seed coat depends heavily on *IQD9*, while cytoskeleton organization in other tissues likely requires additional *IQDs*.

**Fig. 6.**
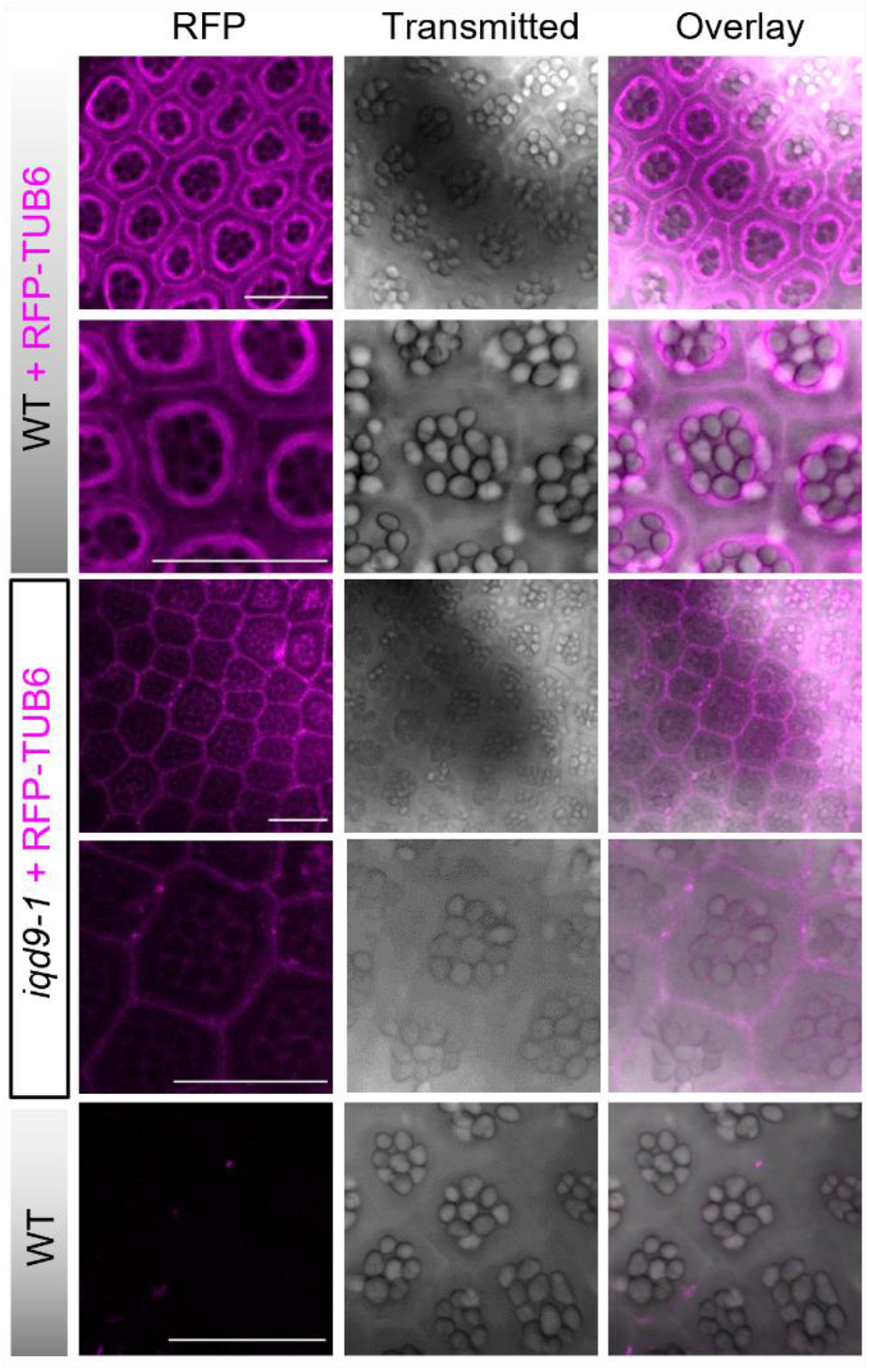
MT organization was impaired in *iqd9* SCE cells. Z-stack maximum projections of RFP-TUB6 in the SCE cells at 7 DPA stage. RFP-TUB6 cortical arrays formed in the WT background but not in *iqd9* SCE cells. WT cells without RFP-TUB6 served as a negative control. Bars = 50 µm.

### The loss of *IQD9* reduces CESA3 velocity

CESA3 is a key subunit of the CSC that polarly deposits cellulose in seed mucilage pockets (Griffiths *et al*., 2015). Since cellulose distribution is disordered in *iqd9* mucilage (Fig. 3), we hypothesized that IQD9 influences CSC motility at the cell cortex. Consistent with previous results (Griffiths *et al*., 2015), time-lapse images revealed GFP-CESA3 proteins moved in a unidirectional, clockwise manner around the cytoplasmic column of SCE cells (Fig. 7). While this pattern was still present, the velocity of GFP-CESA3 particles decreased from 135.9 ± 8.2 nm min^-1^ in WT cells to only 92.8 ± 17.3 nm min^-1^ in *iqd9* (mean ± SD; at least 170 measurements of 9 cells from 3 plants per genotype). Consistent with their mucilage staining phenotypes (Fig. 1, Fig. S2 and Fig. S3), the movement of GFP-CESA3 was also reduced in *trm4* cells, akin to *iqd9* (Fig. 7). In these mutant seeds, CSC movement appeared to be uncoupled from MTs, a behavior previously described for CESA proteins in *klcr* (*cmu*) mutants (Liu *et al*., 2016), but could not be monitored in greater detail due to the severe disruption of RFP-TUB6 localization in SCE cells (Fig. 6; Yang et al., 2019). Taken together, IQD9 is a novel protein required for seed mucilage biosynthesis by maintaining cortical MT arrays and the speed of CESA movement, which influence cellulose distribution.

**Fig. 7.**
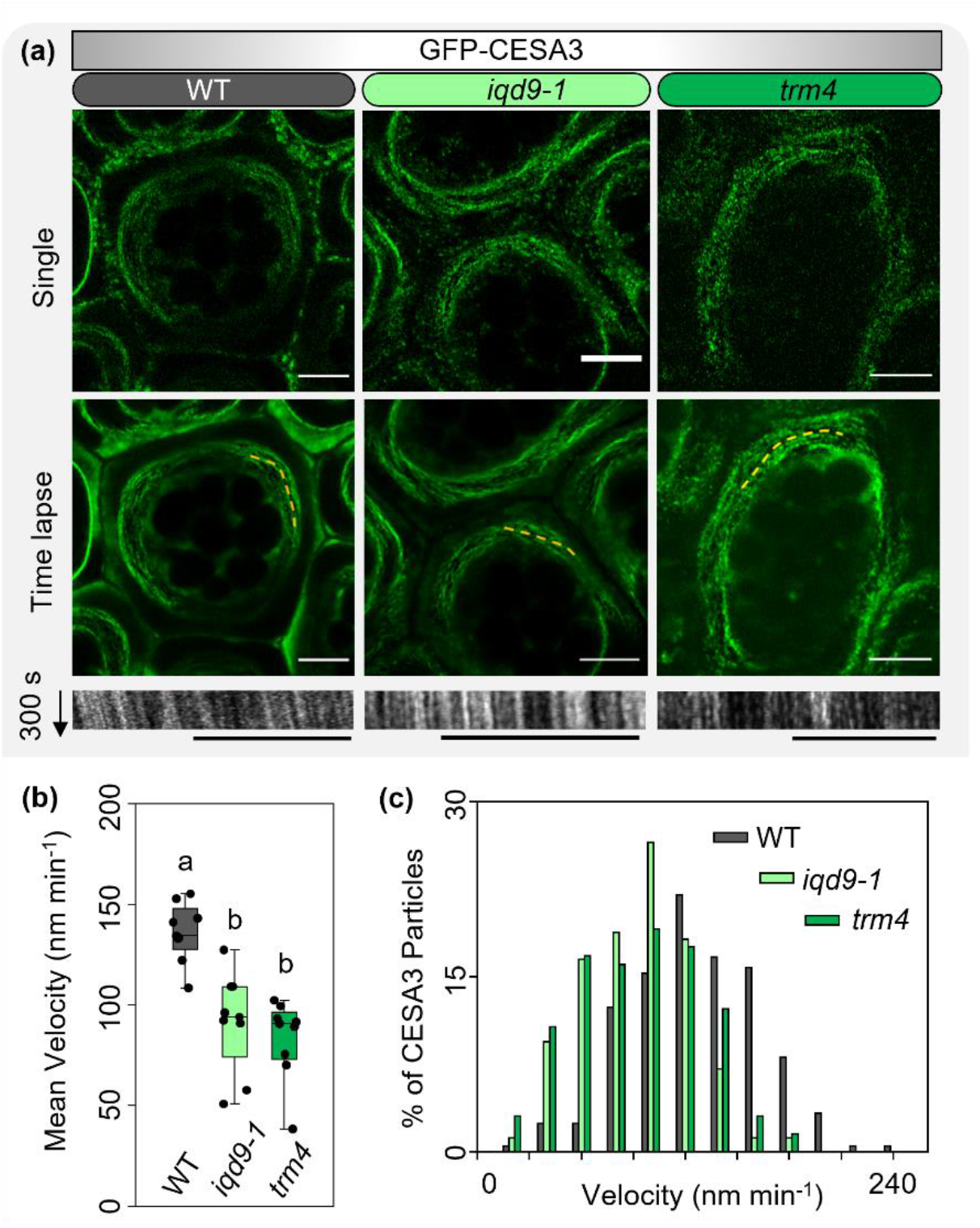
*IQD9* and *TRM4* enhance the velocity of GFP-CESA3 proteins. (a) Single and time-lapse images (acquired every 5 s for 300 s) of GFP-CESA3 in WT, *iqd9* and *trm4* SCE cells at 7 DPA. The bottom row shows kymographs of GFP-CESA3 from dashed lines in the middle row. Bars = 10 µm. (b) Mean GFP-CESA3 velocity in 9 cells from three plants per genotype. Letters label significant differences (one-way ANOVA with Tukey test, P< 0.01). (c) Distribution of GFP-CESA3 velocities for analyzed particles (N=209 for WT, 170 for *iqd9* and 131 for *trm4*).

## Discussion

In the past decade, SCE cells have become a popular model to identify and study cell wall regulators as well as carbohydrate-active enzymes. Dozens of mucilage-related genes have been gradually characterized in Arabidopsis (Voiniciuc *et al*., 2015c; Šola *et al*., 2019), primarily through forward and reverse genetic screens. In addition to mutants generated in the laboratory, the architecture of mucilage β-glucans was found to vary dramatically in natural populations of *Arabidopsis thaliana* (Sullivan *et al*., 2011; North *et al*., 2014; Voiniciuc *et al*., 2016). The Arabidopsis research findings have been accompanied by advances in the mucilage structure of food crops such as *Linum usitatissimum* (flax; Viudes et al., 2020) and *Plantago ovata* (psyllium; (Cowley & Burton, 2021), which contain a higher proportion of non-pectic polymers. Despite the evolution of various mucilage traits within the *Brassicaceae* family (Viudes *et al*., 2021), how MTs regulate the intricate organization of this specialized secondary cell wall has remained a relatively blank slate. The Arabidopsis genome encodes hundreds of putative MT-associated proteins, but only MOR1 and TRM4 were previously shown to influence seed mucilage synthesis (McFarlane *et al*., 2008; Hamada, 2014; Yang *et al*., 2019). Additional MT-associated proteins (e.g. CSI1/POM2; CC1; IQD13 and KLCRs/CMUs) involved in cell wall biosynthesis were characterized in other tissues (Li *et al*., 2012; Bringmann *et al*., 2012b; Endler *et al*., 2015; Liu *et al*., 2016; Sugiyama *et al*., 2017), so the players that guide mucilage biosynthesis remained unclear.

### IQD9 sustains MT organization during specialized cell wall deposition

In this study, we discovered that IQD9 and its interactor KLCR1 localize to cortical arrays that resemble the circular paths of MTs (Fig. 6) and multiple CESAs (Griffiths *et al*., 2015) during mucilage biosynthesis. IQD9 co-localized with MTs and was sensitive to their depolymerization by oryzalin, suggesting that IQD9 may be capable of directly binding MTs like other family members. The DUF4005 domain of IQD16 was recently shown to mediate MT binding *in vivo* as well as *in vitro* (Li *et al*., 2021). As one of the shortest family members (Abel *et al*., 2005), IQD9 lacks the DUF4005 domain but contains a region similar to the MT2 domain of IQD13 (Fig. S1c), which is sufficient for MT localization *in vivo* (Sugiyama *et al*., 2017). The highly immobile IQD9 proteins could function similarly to KLCR1/CMU1, its binding partner (Fig. 4), to stabilize the cortical MT arrays of SCE cells and sustain CSC speed during cellulose deposition. Consistent with this hypothesis, oryzalin treatment of SCE cells severely disrupted the trajectory and velocity of GFP-CESA3 (Griffiths *et al*., 2015). The reduced velocity of CESA3-containing CSCs in *iqd9* and *trm4* SCE cells (Fig. 7) shows that multiple classes of proteins are required to shape the circular MT arrays and cellulose distribution.

### MT organization primarily affects cellulose distribution in seed mucilage

In Arabidopsis, SCE cells display MT and CSC dynamics that are considerably different from those of hypocotyl and protoxylem vessels (Griffiths *et al*., 2015; Watanabe *et al*., 2015; Griffiths & North, 2017), which serve as primary and secondary cell wall models. Both hypocotyl or protoxylem cells show transverse MTs arrays or bundles, aligned with bidirectional movement of CSCs in the PM. The velocities of CSCs during cellulose deposition range from 200–300 nm min^-1^ in the hypocotyl to 300–400 nm min^-1^ in the protoxylem. By contrast, SCE cells display circular MT arrays around the columella, aligned with unidirectional movement of CSCs with a velocity of 80 to 120 nm min^-1^ (Fig. 7, Griffiths et al., 2015). These unique MT pattern and CSC movements could lead to the polarized deposition of unusual cellulosic coils, which unwind with the expansion of pectin polymers during Arabidopsis seed hydration (Šola *et al*., 2019). Cytoskeletal defects in *iqd9, klcr1*, or *trm4* mutants could slow CSC movement to result in shorter cellulose microfibrils that cannot extend to form long ray-like structures during mucilage release. The compact mucilage phenotypes of MT-related mutants and galactoglucomannan-deficient seeds such as *muci10* could be explained by similar deficiencies in the assembly of cellulose chains (Yu *et al*., 2014; Voiniciuc *et al*., 2015b; Griffiths & North, 2017; Yang *et al*., 2019), Since relatively abundant MTs line the PM of the mucilage pocket where mucilage secretion occurs (McFarlane *et al*., 2008), MTs could also potentially target the secretion of pectin and hemicelluloses to the apoplast. However, a temperature-sensitive mutation of *MOR1* partially disrupted mucilage release without clearly affecting the secretion of vesicles to the mucilage pocket and mucilage polymer accumulation (McFarlane *et al*., 2008). Despite severely disrupted RFP-TUB6 localization in SCE cells (Fig. 6; Yang et al., 2019), the *iqd9* and *trm4* seeds released matrix polysaccharides with a composition that was similar to WT (Fig. 2). Even though the *iqd10-1* mutation partially reduced certain monosaccharides (Fig. 2), the mucilage released from these seeds showed a WT-like appearance (Figs. 1 and 3). Therefore, cortical MTs appear to have a relatively minor impact on the incorporation of pectin and hemicelluloses into the mucilage pockets of SCE cells. Unaltered mucilage adherence to the surface of *iqd9, klcr1* and *trm4* seeds is likely mediated by the presence of xylan (Voiniciuc *et al*., 2015a; Ralet *et al*., 2016), the SOS5 arabinogalactan protein and the receptor-like kinase FEI2 (Harpaz-Saad *et al*., 2011; Griffiths *et al*., 2016; Šola *et al*., 2019).

### IQD, KLCR, and TRM proteins have interconnected functions

Our results and recently published findings indicate that distinct IQD proteins function in a tissue-specific manner by interacting with KLCR proteins and have potentially overlapping roles with TRM scaffolding proteins. Based on double mutant analyses (Figs. S2 and S3), the roles of *IQD9* in SCE cells are nearly identical to those of *KLCR1* and *TRM4*. Therefore, the encoded proteins could be associated as part of a single complex or pathway at the cell cortex. Some Arabidopsis IQDs (GFP-IQD1 or GFP-IQD2) can recruit KLCR1 to MTs when transiently co-expressed in tobacco cells (Bürstenbinder *et al*., 2013; Zang *et al*., 2021). The MT recruitment is consistent with our results for the transient expression of IQD9-GFP and KLCR1 (which behaves similarly when tagged with either RFP or mCherry), even though KLCR1/CMU1 alone can at least partially bind MTs (Liu *et al*., 2016; Zang *et al*., 2021). Furthermore, the KLCR1-IQD2 pair was recently shown to interact with the actin binding protein NET3C to modulate the shape of the endoplasmic reticulum at PM contact sites (Zang *et al*., 2021). We hypothesize that IQD9, KLCR1, TRM4 could be an integral part of an expanding group of proteins (Polko & Kieber, 2019) that support PM-bound CSC movement along the orientation of MT tracks. IQDs are hypothesized to function as scaffolds and may be modified by TON1/TRM/PP2A (TTP)-mediated dephosphorylation (Kumari *et al*., 2021). Via IQ67-domain-mediated calmodulin binding (Abel *et al*., 2005), IQDs could also participate in Ca^2+^ signaling to ultimately influence plant cell wall dynamics.

### Intriguing roles of IQDs during secondary cell wall formation

Even though *IQD9* and *IQD10* promoters were active in both vegetative and reproductive organs (Fig. S1), *IQD9* was indispensable only for the organization of seed mucilage polysaccharides (Figs. 1–3). While the expression of IQD9-GFP under its native promoter was detected only in the general seed coat (Fig. 5 and Fig. S4), KLCR1-GFP was expressed more ubiquitously in complemented lines. The transcription of *IQD9, IQD10* and *IQD13* was previously associated with secondary cell wall biosynthesis (Mutwil *et al*., 2008). Even though we detected transcriptional activity in the vasculature (Fig. S1), the absence of *IQD9* and/or *IQD10* did not cause *irx* phenotypes found in stems with defective cellulose-hemicellulose networks (Fig. S6; Brown et al., 2007). Their vascular functions could be masked by the expression of related genes such as *IQD13*, which was already shown to modulate MT organization during xylem cell formation (Sugiyama *et al*., 2017). In poplar, the down-regulation of *PdIQD10* during wood formation increased the tree height, diameter and relative cellulose content (Badmi *et al*., 2018). Since PdIQD10 interacted with PdKLCRs and could be directed to the nucleus, the elevated cellulosic biomass of the transgenic trees suggests that IQD-KLCRs participate in a tight feedback-loop that regulates cellulose biosynthesis (Badmi *et al*., 2018). In Arabidopsis, multiple *IQDs* likely have redundant roles during stem development so higher-order mutants would be needed to decipher how IQD-KLCR are involved in signaling pathways or in direct interactions with secondary wall CSCs.

### Future avenues to tailor cellulose deposition

The localization of soluble IQD9 and KLCR1/CMU1 (Fig. 5, Liu et al., 2016) proteins near or at the PM suggests that they could interact with membrane-bound CSC components, which travel in a spiral pattern during mucilage synthesis (Griffiths *et al*., 2015). TRM4 was previously shown to maintain MT organization and directly bind CESA3 (Yang *et al*., 2019) to enhance its mobility (Fig. 7). Although the mechanism that connects IQD9 and KLCR1 to TRM4 requires further investigation, we provide the first evidence that members of these three MT-associated families cooperate to direct cellulose deposition. Additional CSC-related genes are expressed during mucilage production (Griffiths & North, 2017), but their putative roles in mucilage biosynthesis remain to be investigated. Exploring the interactome of IQD9, KLCR1, and TRM4 could reveal novel targets to fine-tune the biosynthesis of cellulose, the most abundant renewable material on our planet. In addition to plant studies, the growing arsenal of proteins found to influence cellulose biosynthesis could be rapidly expressed and engineered in surrogate hosts (Pauly *et al*., 2019). Yeast species such as *Pichia pastoris* have already been used to express a *Populus* CESA capable of producing cellulose microfibrils *in vitro* (Purushotham *et al*., 2016) and to identify essential protein co-factors for CESA-like enzymes that catalyze hemicellulose elongation (Voiniciuc *et al*., 2019). Therefore, synthetic biology advances combined with attractive plant models, such as the Arabidopsis SCE cells, provide exciting avenues to refine the fibers that shape plants and many industrial products.

## Acknowledgements

We thank Tilman Jacob, Romina Plötner and Kristina Rosenzweig for technical assistance and colleagues at the Leibniz Institute of Plant Biochemistry for microscope access. The *pUBQ:RFP-TUB6, pKLCR1:KLCR1-GFP* in *klcr1-1* seeds and the *proCESA3:GFP-CESA3* in *je5* seeds were provided by Geoffrey Wasteneys (University of British Columbia, CA), Pengwei Wang (Huazhong Agricultural University, CN) and Samantha Vernhettes (Université Paris-Saclay, FR), respectively. This work was supported by core funding (Leibniz Association) from the Federal Republic of Germany and the state of Saxony-Anhalt, and by DFG grants (BU2955/2-1 and BU2955/1-1 to K.B.; 414353267 to C.V.).

## Authors Contributions

C.V. and K.B. conceived the project. B.Y. performed most of the experiments under C.V.’s supervision. G.S. and K.B. contributed genetic materials and performed RT-PCR, GUS and co-IP assays. B.Y. and C.V. analyzed data and wrote the manuscript with input from all authors.

