## Supporting Information for "Microtubule-associated IQD9 guides cellulose synthase velocity to shape seed mucilage"

The following Supporting Information is available for this article:

**Fig. S1** *IQD9* and *IQD10* promoter activities and protein sequence alignment.

**Fig. S2** *IQD9* influences mucilage deposition like *KLCR1* and *TRM4*.

**Fig. S3** Mucilage β-glucan staining of *iqd9*, *klcr1* and *trm4* mutants.

**Fig. S4** SCE cells visualized with propidium iodide staining.

**Fig S5** *iqd9* and *klcr1* are not sensitive to salt stress.

**Fig. S6** Transverse sections and monosaccharide composition of stems.

**Fig. S7** IQD9 co-aligns with cortical microtubules.

**Fig. S8** KLCR1 and IQD9 localization during SCE cell development.

**Fig. S9** RFP-TUB6 localizes to cortical arrays in *iqd9* hypocotyl.

**Table S1** Mutants used in this study.

**Table S2** Primers used in this study.

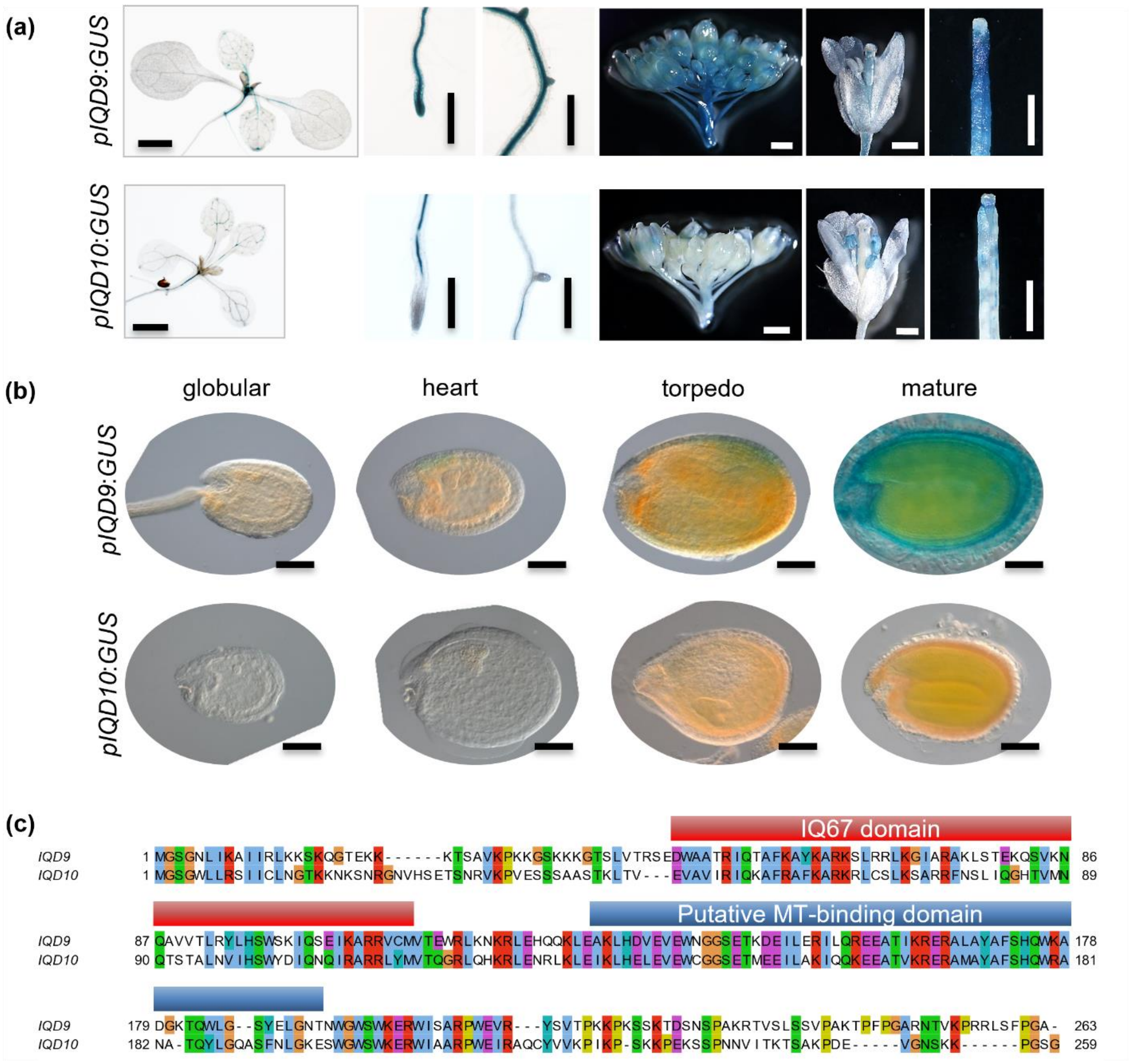


**Fig. S1** *IQD9* and *IQD10* promoter activities and protein sequence alignment.
(a) Promoter:GUS activities were examined in shoots of 10-day-old seedling, as well as in the flower buds, mature flowers, and siliques of 6–7-week-old plants. (b) GUS activity in four stages of developing seeds. Bars = 1 mm in (a) and 100 µm in (b). (c) MUSCLE sequence alignment of IQD9 and IQD10 visualized using Jalview (CulstalX colors for different amino acids). IQD9 is proposed to bind CaM via the eponymous IQ67 domain (Abel et al., 2005). A putative MT-binding region was annotated based on similarity to the MT2 domain of IQD13 (Sugiyama et al., 2017).

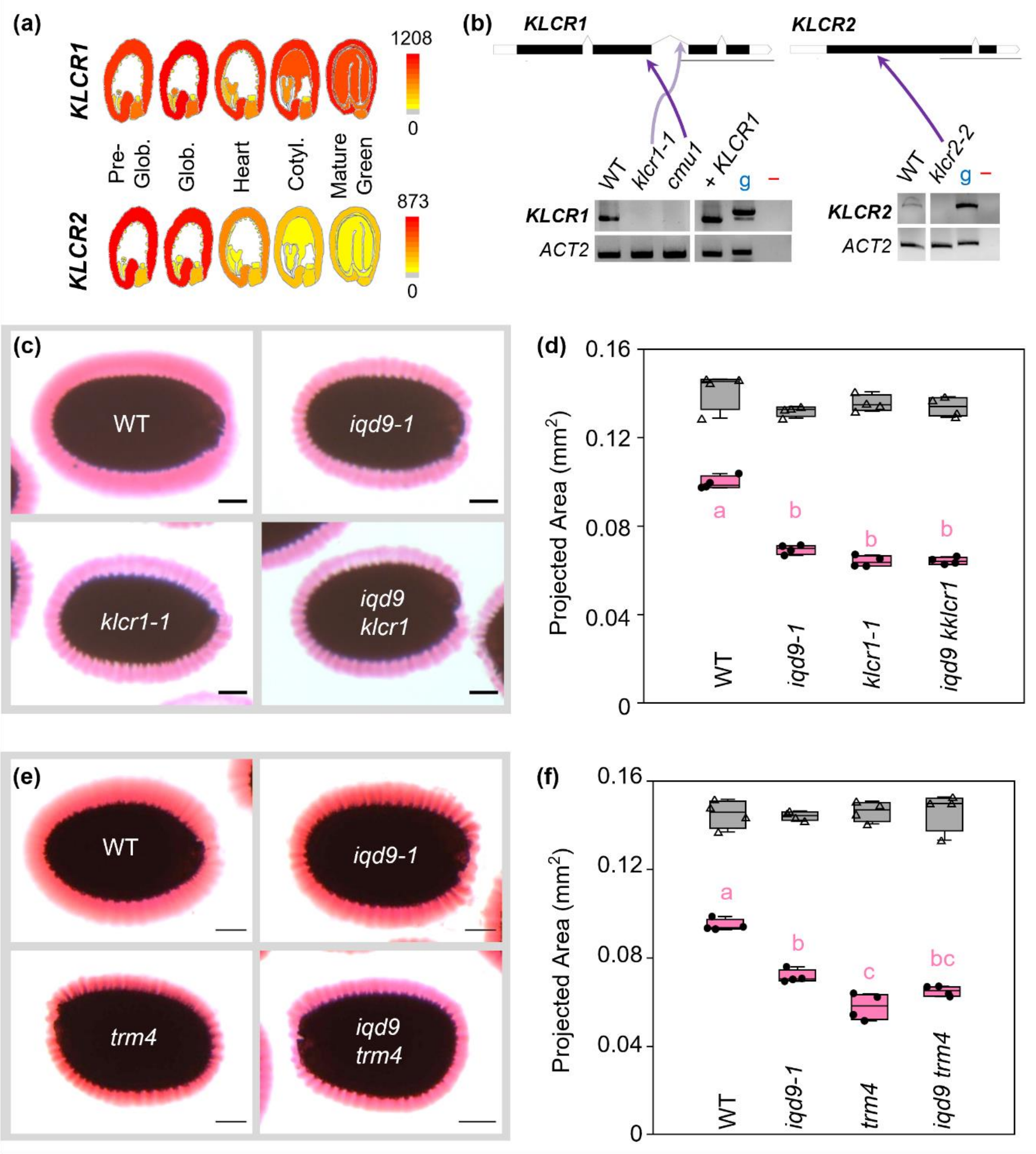


**Fig. S2** *IQD9* influences mucilage deposition like *KLCR1* and *TRM4*.

(a) Expression profiles of *KLCR1* and *KLCR2* in the seed eFP browser (Winter *et al*., 2007; Le *et al.*, 2010), including absolute expression values. Glob. (globular); Cotyl. (cotyledon). (b) UTR, intron and exon structure of KLCR genes. The position and the effects of T-DNA insertions were verified using RT-PCR, *ACTIN2* as a reference gene, g as genomic DNA control, and – as no DNA control. Bars for gene models = 1000 bp. (c) and (e) Representative images of RR-stained seeds. Bars = 100 µm. (d) and (f) Areas of seeds (triangles) and adherent mucilage (dots). Boxes show the 25–75% quartiles, the median value (inner horizontal line), and whiskers extending to the largest/smallest values. Data are from four biological replicates (≥20 seeds each). Different letters mark significant changes (one-way ANOVA with Tukey test, P < 0.01).

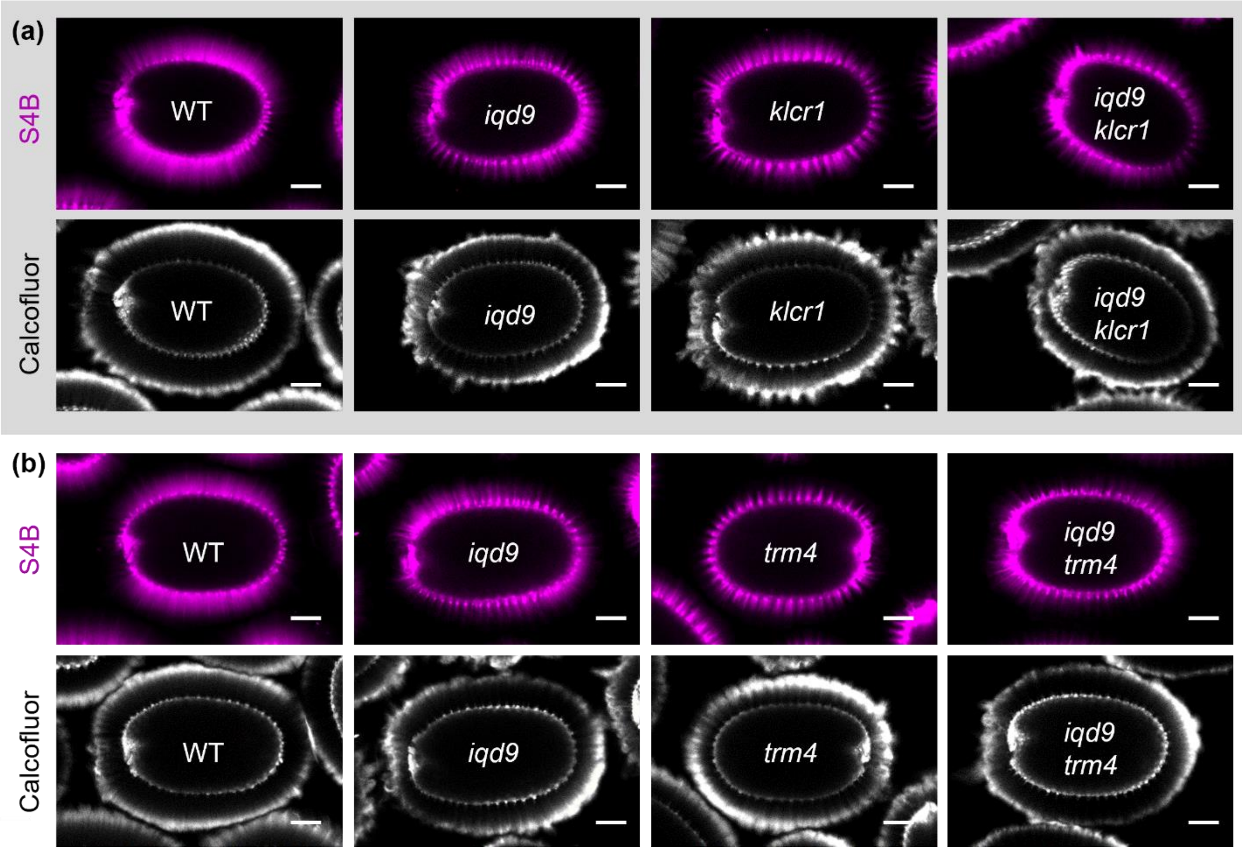


**Fig. S3** Mucilage β-glucan staining of *iqd9*, *klcr1* and *trm4* mutants.
(a) and (b) Cellulosic rays were labelled with S4B, and the mucilage capsule was then counterstained with Calcofluor White, a β-glucan dye with broader specificity. Bars = 100 µm.

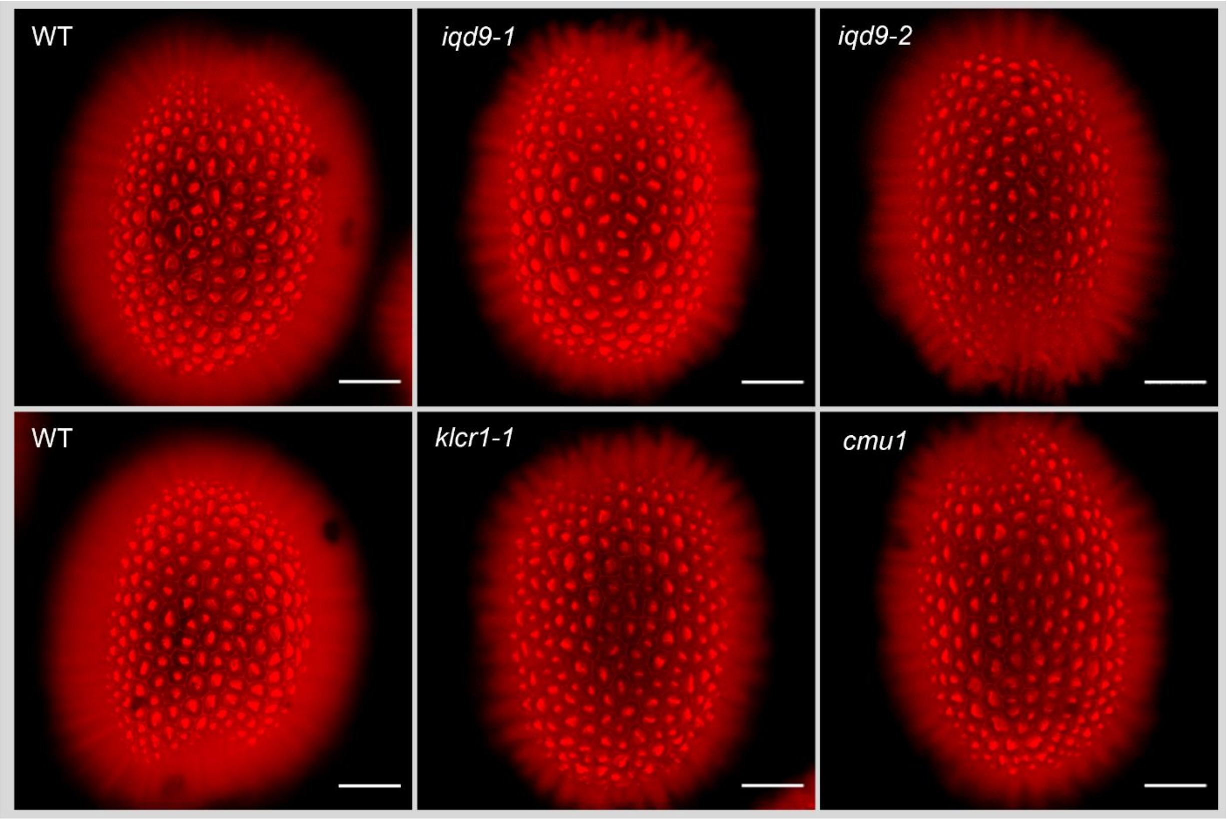


**Fig. S4** SCE cells visualized with propidium iodide staining.
Representative images of seeds showing epidermal cell morphology. The sizes and shapes of SCE cells were indistinguishable for these genotypes. Bars = 100 µm.

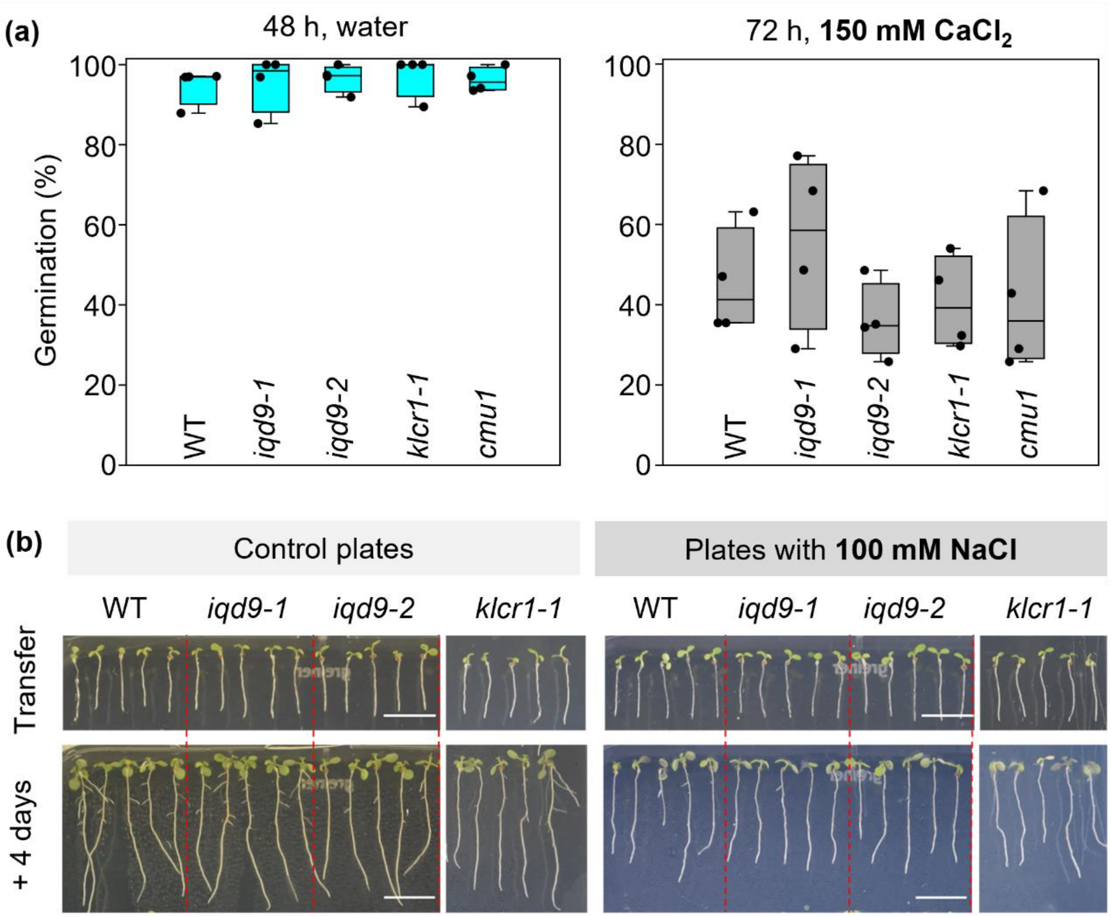


**Fig. S5** *iqd9* and *klcr1* are not sensitive to salt stress.

(a) Seed germination rate in water and 150 mM $CaCl_2$. (b) 5-day-old seedlings were transferred to Arabidopsis agar media plates with or without 100 mM NaCl. The upper panel shows the seedlings immediately upon transfer. The lower panel displayed the seedlings after an additional 4 days of growth. Root elongation was impaired after 4 days salt stress treatment, but no major differences were observed between the genotypes. Bars = 1 cm.

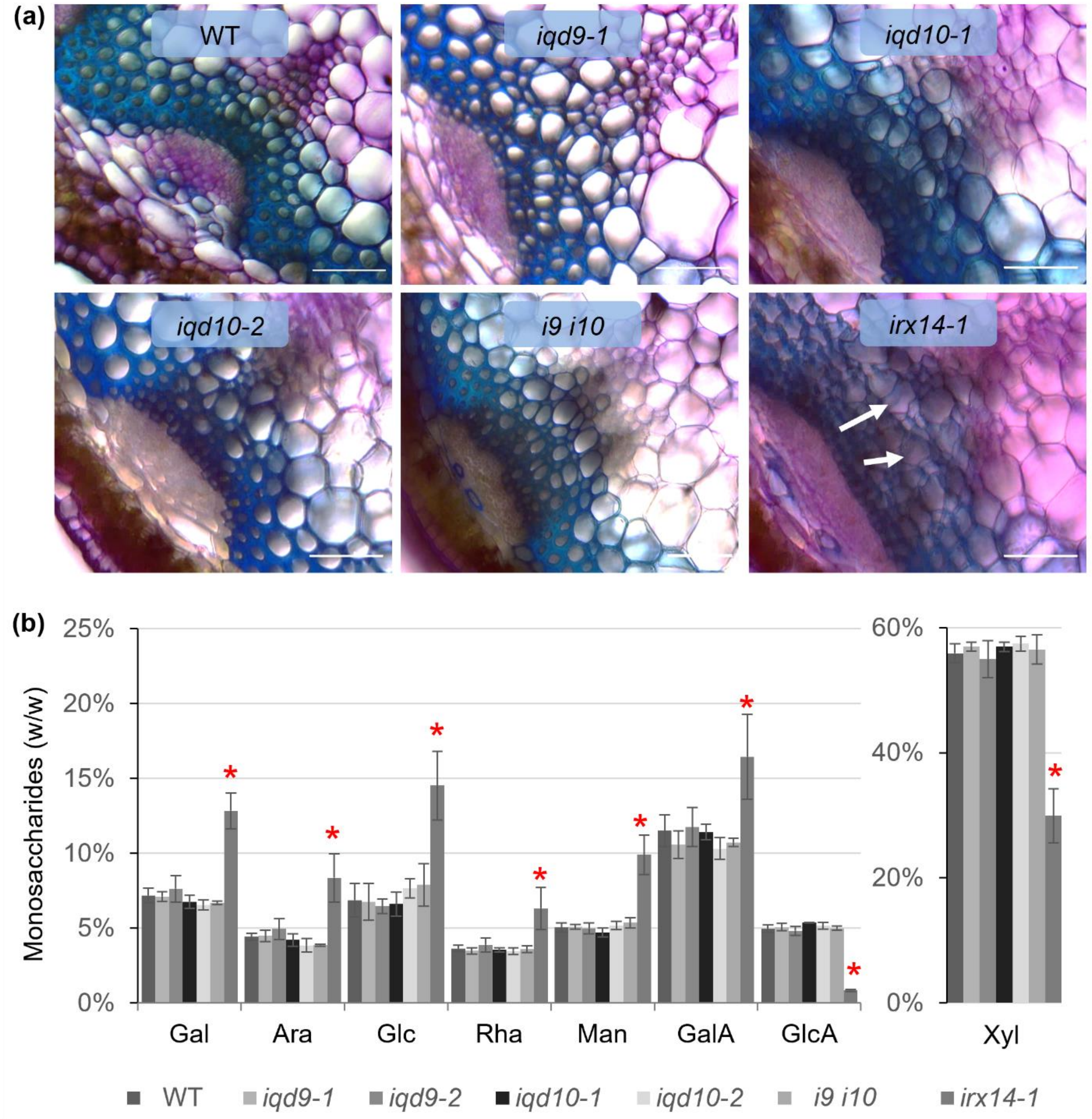


**Fig. S6** Transverse sections and monosaccharide composition of stems.
(a) Transverse sections of Arabidopsis stems stained with toluidine blue O. Arrows indicate collapsed xylem elements. The brightness of each subpanel was adjusted independently to yield a comparable view of xylem cells. Bars = 50 µm. (b) Monosaccharide composition of the mature stems. Values represent means ± SD of five biological replicates. The asterisk marks significant decreases compared with WT (Student's t-test, P < 0.05).

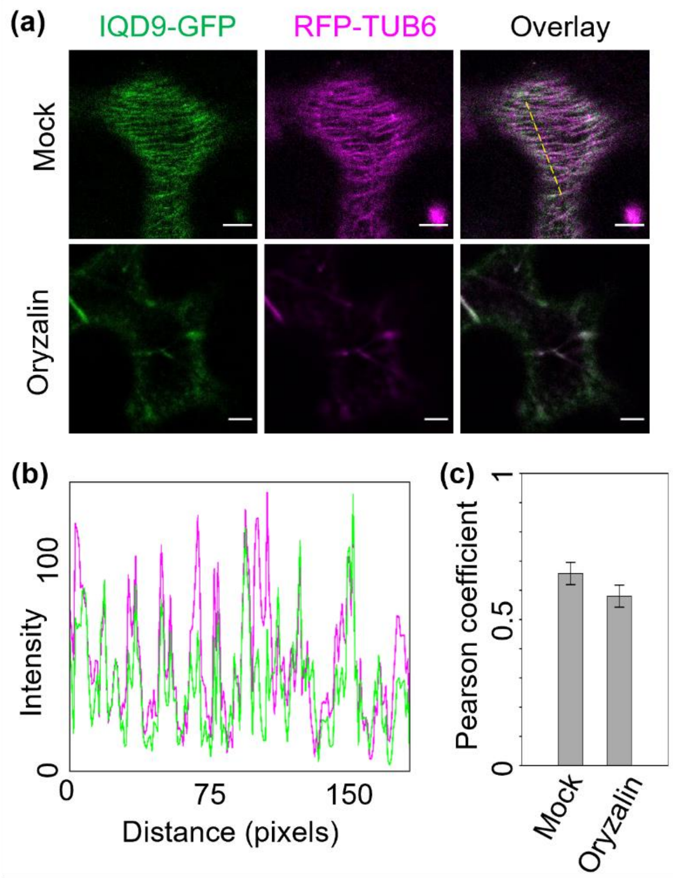


**Fig. S7** IQD9 co-aligns with cortical microtubules.
(a) Mock and oryzalin treatment of IQD9-GFP and RFP-TUB6 transient expressed in *N. benthamiana* leaf epidermal cells. Both IQD9-GFP and RFP-TUB6 are sensitive to oryzalin treatment. (b) Fluorescent intensity plot along the dashed line in (a). (c) Pearson correlation coefficient between the IQD9-GFP and RFP-TUB6 in the mock and oryzalin treatment, n=5 cells from 5 independent treatments. Bars = 10 μm.

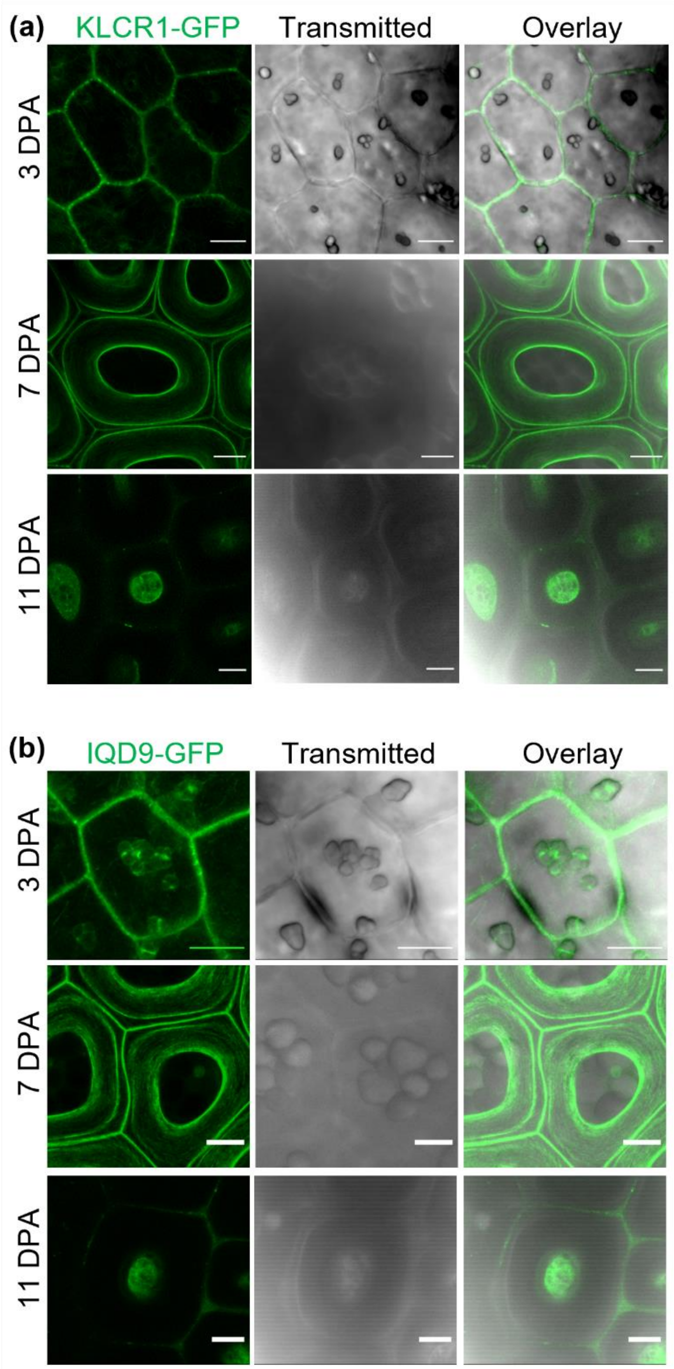


**Fig. S8** KLCR1 and IQD9 localization during SCE cell development.
(a) Maximum Z-stack projections of *klcr1-1* + *KLCR1-GFP* seeds at 3, 7 and 11 DPA. Starch granules, a mucilage donut-shaped pocket, and the columella were prominent at 3, 7 and 11 DPA, respectively. (b) Maximum Z-stack projections of *iqd9-1* + *IQD9-GFP* seeds at three developmental stages, which were also analyzed for KLCR1-GFP in (a). Bars = 10 µm.

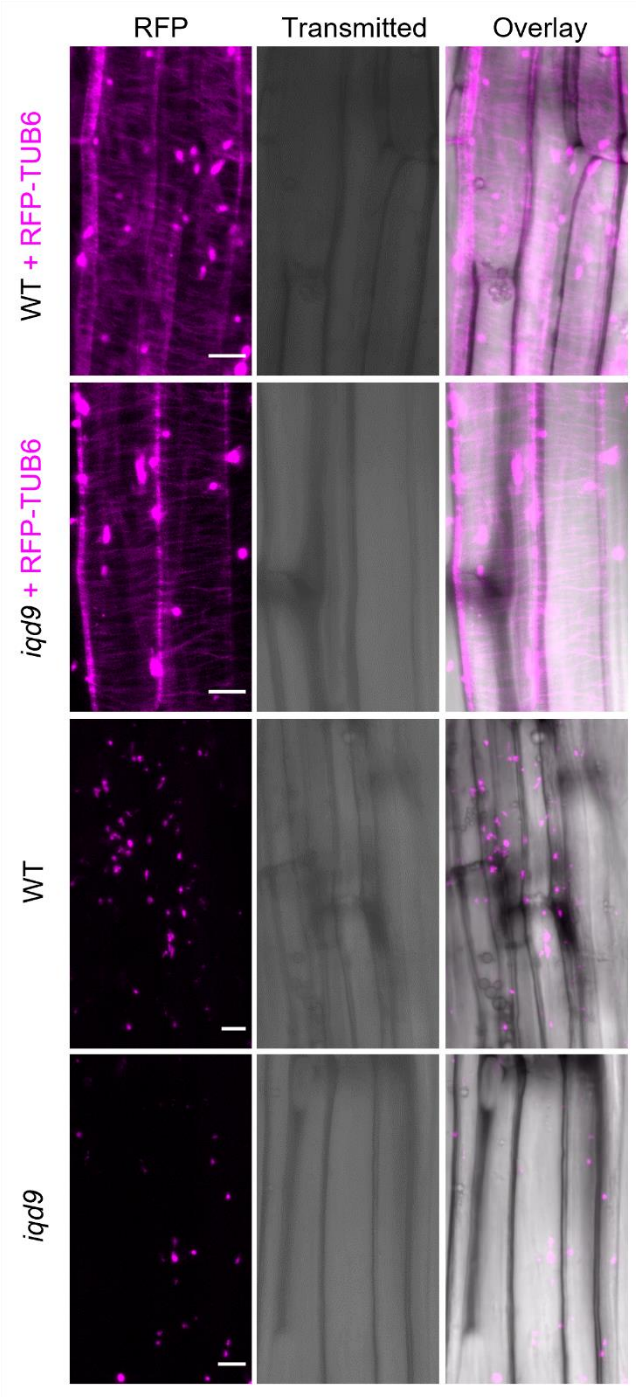


**Fig. S9** RFP-TUB6 localizes to cortical arrays in *iqd9* hypocotyl.
Maximum projection of Z-stack images of RFP-TUB6 on the inner face in the zone 1 of hypocotyl of WT and *iqd9* lines that were used for seed coat imaging. WT and *iqd9* hypocotyl cells without the MT marker only showed chloroplast intrinsic fluorescence. Bars = 10 µm.

**Table S1** Mutants used in this study.
The references indicate where the alleles were first described.

| Mutant | Locus | Polymorphism | Reference |
|---|---|---|---|
| *iqd9-1* | At2g33990 | SALK_041348 | This study |
| *iqd9-2* | At2g33990 | WiscDsLox287H03 | This study |
| *iqd10-1* | At3g15050 | SALK_111401 | This study |
| *iqd10-2* | At3g15050 | SAIL_266_A09 | This study |
| *klcr1-1* | At4g10840 | SAIL_335_B08 | (Ganguly et al., 2020) |
| *cmu1* | At4g10840 | SAIL_273_F11 | (Liu et al., 2016) |
| *klcr2-2* | At3g27960 | SALK_148296 | (Liu et al., 2016) |
| *trm4-1* | At1g74160 | SALK_022851 | (Yang et al., 2019) |
| *muci10-1* | At2g22900 | SALK_061576C | (Voiniciuc *et al.*, 2015) |
| *irx14-1* | At4g36890 | SALK_038212 | (Brown *et al.*, 2007) |

**Table S2** Primers used in this study.
The primers were sorted by primary application (genotyping, RT-PCR and cloning), and the target sequence specified in the description. The table is continued on the next page.

| | Name | Primer sequence (5\`-> 3\`) | Description |
|---|---|---|---|
| Plant Genotyping | BY012F | ATTTTGCCGATTTCGGAAC | SALK LBb1.3 |
| | BY013F | TAGCATCTGAATTTCATAACCAATCTCGATACAC | SAIL LB3 |
| | BY015F | AACGTCCGCAATGTGTTATTAAGTTGTC | Wisc LB p745 |
| | BY060F | TTTGTCTCTGACCCACCATTC | *iqd9-1* |
| | BY060R | TTCATTGGTCCTGAACTCAGG | |
| | BY067F | AACCACTAAATTCGATTCCGG | *iqd9-2* |
| | BY067R | GTCCATCACATTTTGTGCAAG | |
| | BY061F | TGTGTAGCATTAGCCCTCCAC | *iqd10-1* |
| | BY061R | GCTCAGTTGGTTAGAGCGTTG | |
| | BY062F | AACAGCACCACCTTGAATTTG | *iqd10-2* |
| | BY062R | TGATCGGTTTAACGACGTAGC | |
| | BY064F | GATCAATCAGATTCTGGAGCG | *klcr1-1* |
| | BY064R | TATACAATAAGCCGGTGCCTG | |
| | BY084R | GGACTTTACCTTCCCATAGCG | *cmu1* |
| | BY084F | GCGAGTGGACAAGAATCTGAG | |

| | Name | Primer sequence (5`-> 3`) | Description |
|---|---|---|---|
| | BY065F | GTTCCTGGAGTCGGTTTTAGG | *klcr2-2* |
| | BY065R | GTCCAGGGCGAGTTTTATTTC | |
| | BY066F | ATGGTTCGTGTCCAGTAGTCG | *trm4-1* |
| | BY066R | TGGGATCAACACTGGAGTTTC | |
| RT-PCR | BY012F | CACC-ATGGGTTCTGGGAATTTGATT | IQD9 |
| | IQD125 | TCAAGCACCTGGAAATGACA | |
| | IQD126 | CACC-ATGGGATCTGGATGGCTG | IQD10 |
| | IQD127 | TTATCCGGAACCAGGCTTT | |
| | IQD300 | CACC-ATGCCAGCAATGCCAGGT | KLCR1 |
| | IQD301 | TCAGAACTTGAAACCGAGGC | |
| | IQD326 | CACC-ATGGACGTAGGAGAGAGCAATG | KLCR2 |
| | IQD327 | TCAATAAACCGGTCTCTGTCC | |
| Cloning primer | IQD188 | CACC-GCTAATTAACACACACAAAATTTAGAA | ProIQD9 amplification |
| | IQD189 | CTTAGAGTCTATCAATTGTCACCAAAA | |
| | IQD190 | CACC-TTTTTGAGACTTTCAGGTCACC | ProIQD10 amplification |
| | IQD191 | AACAGAAGACTGGCTCAGTTTAAAG | |
| | IQD124 | CACC-ATGGGTTCTGGGAATTTGATT | IQD9 CDS |
| | IQD125 | TCAAGCACCTGGAAATGACA | |
| | IQD300 | CACC-ATGCCAGCAATGCCAGGT | KLCR1 CDS |
| | IQD301 | TCAGAACTTGAAACCGAGGC | |
| | IQD1159 | GGGGACAAGTTTGTACAAAAAAGCAGGCTTCATGGAATCCGAAGG AGAAAC | CESA3 CDS |
| | IQD1384 | GGGGACCACTTTGTACAAGAAAGCTGGGTCTCACAACAGTTGATTC CACATT | |
| | IQD1607 | GGGGACAAGTTTGTACAAAAAAGCAGGCTTCGCTAATTAACACAC ACAAAATTTAGAA | IQD9 gDNA |
| | IQD1608 | GGGGACCACTTTGTACAAGAAAGCTGGGTCAGCACCTGGAAATGA CA | |
